# Epidermal Langerhans Cells Act as Neuroimmune Regulators of Pain and Itch

**DOI:** 10.64898/2026.11.06.756111

**Authors:** Kent Sakai, Hirotake Ishida, Darya Pavlenko, Masataka Suehiro, Thomas Oshea, Jessica Dorning, Kristen Sanders, Charel Sanchez, Ariana Lopez, Sophia Talamas, Maria Boulina, Anthony J Griswold, Tasuku Akiyama

## Abstract

Neuroimmune interactions in the skin coordinate immune defense and somatosensory perception, yet how immune cells shape sensory signaling remains poorly defined. Here, we show that Langerhans cells (LCs), epidermal antigen-presenting cells, function as local immune-derived amplifiers of cutaneous sensory signaling. Anatomical and transcriptomic analyses reveal preferential LC association with non-peptidergic sensory neurons. LC activation increases neuronal and behavioral responses to thermal, mechanical, and pruritic stimuli without directly exciting sensory neurons, whereas LC inhibition or ablation selectively reduces pruritogen-evoked itch. We identified semaphorin 7A as a candidate LC-expressed mediator. Semaphorin 7A sensitized cultured sensory neurons to chloroquine and histamine, and neutralizing semaphorin 7A abolished the increase in chloroquine-evoked scratching produced by LC activation. Its receptor PlexinC1 was expressed by most MrgprA3⁺ pruriceptors and blocking PlexinC1 reduced semaphorin 7A-evoked neuronal responses. Together, these findings identify epidermal LCs as local immune-derived amplifiers that regulate the responsiveness of cutaneous sensory circuits and identify semaphorin 7A as a mediator of LC-induced itch potentiation.

## INTRODUCTION

Langerhans cells (LCs), first reported by Paul Langerhans in 1868, are specialized antigen-presenting cells that reside in the epidermis (*1, 2*). While they were initially classified within the broader family of antigen-presenting cells, research later established LCs as a special immune population uniquely adapted to the skin (*3, 4*). Located within the epidermal layer, LCs extend their dendritic projections throughout the epidermis, constantly monitoring the external environment and creating a dense network for immune defense (*3, 4*). This allows them to detect pathogens efficiently while maintaining the skin’s barrier, making LCs essential regulators of immune surveillance in the skin.

Increasing evidence indicates that LCs, like other immune cells, are functionally modulated by sensory neurons (*5, 6*). Sensory afferents establish close contact with LCs via axon-like structures, and the neuropeptides such as calcitonin gene-related peptide (CGRP), substance P, pituitary adenylate cyclase-activating peptide, and vasoactive intestinal polypeptide, released from sensory terminals, modulate LC-mediated antigen presentation (*7–9*). Conversely, LCs contribute to the homeostasis of sensory neurons, including both peptidergic and non-peptidergic subtypes (*10–12*). Although these neuronal populations are well established as mediators of pain and itch (*13*), whether and how LCs actively shape sensory processing remains incompletely understood. Here, we combined anatomical, transcriptomic, optogenetic, chemogenetic, and pharmacological approaches to define how LCs interact with sensory neurons and modulate pain and itch.

## RESULTS

### LCs preferentially associate with non-peptidergic sensory fibers

To characterize the neuronal types of sensory neurons closely associated with LCs, we performed immunofluorescent staining on both hairy and glabrous skin. By crossing the Lang^CreERT2^ mice with Ai14 reporter mice, we generated mice in which LCs express tdTomato (**Fig. 1A**). After tamoxifen injections, the Lang^CreERT2^; Ai14 reporter mice showed robust expression of tdTomato in the LCs immunostained with CD207, a marker for LCs (tdTomato/CD207 = 92.9% (289/311), CD207/tdTomato = 98.3% (289/294); *n* = 2-3 sections/mouse from 4 mice; **Fig. 1B**). These data confirm high recombination efficiency and specificity of the reporter system. To characterize the spatial relationship between LCs and cutaneous nerve fibers, we analyzed consecutive 1 µm optical sections through the epidermis (**Fig. S1**). Close proximity was defined as direct overlap between LC profiles and immunolabeled fiber profiles within a given optical section. LC cell bodies exhibited preferential association to GFRα2-positive non-peptidergic C-fibers 64.5% (149/231), compared with more limited associations with CGRP-expressing peptidergic fibers 27.0% (54/200) and neurofilament 200-positive myelinated fibers (22.4%, 30/134) (*n* = 3-5 sections/mouse from 5-6 mice; **Fig. 1C-G**). LC dendrites showed an even stronger bias toward non-peptidergic fibers, contacting the majority of GFRα2-positive axons (78.4%, 181/231), while exhibiting moderate associations with peptidergic fibers (29.5%, 59/200) and sparse interactions with myelinated fibers (10.4%, 14/134). Consistent with this broad innervation, pan-neuronal βIII-tubulin-positive fibers were in close proximity to most LC cell bodies (80.0%, 152/190) and dendrites (79.5%, 151/190) (*n* = 3-4 sections/mouse from 5 mice), indicating that individual LCs reside within dense cutaneous nerve networks. Similar patterns of neuronal contact were observed in non-hairy skin (**Fig. S2**).

**Fig. 1.**
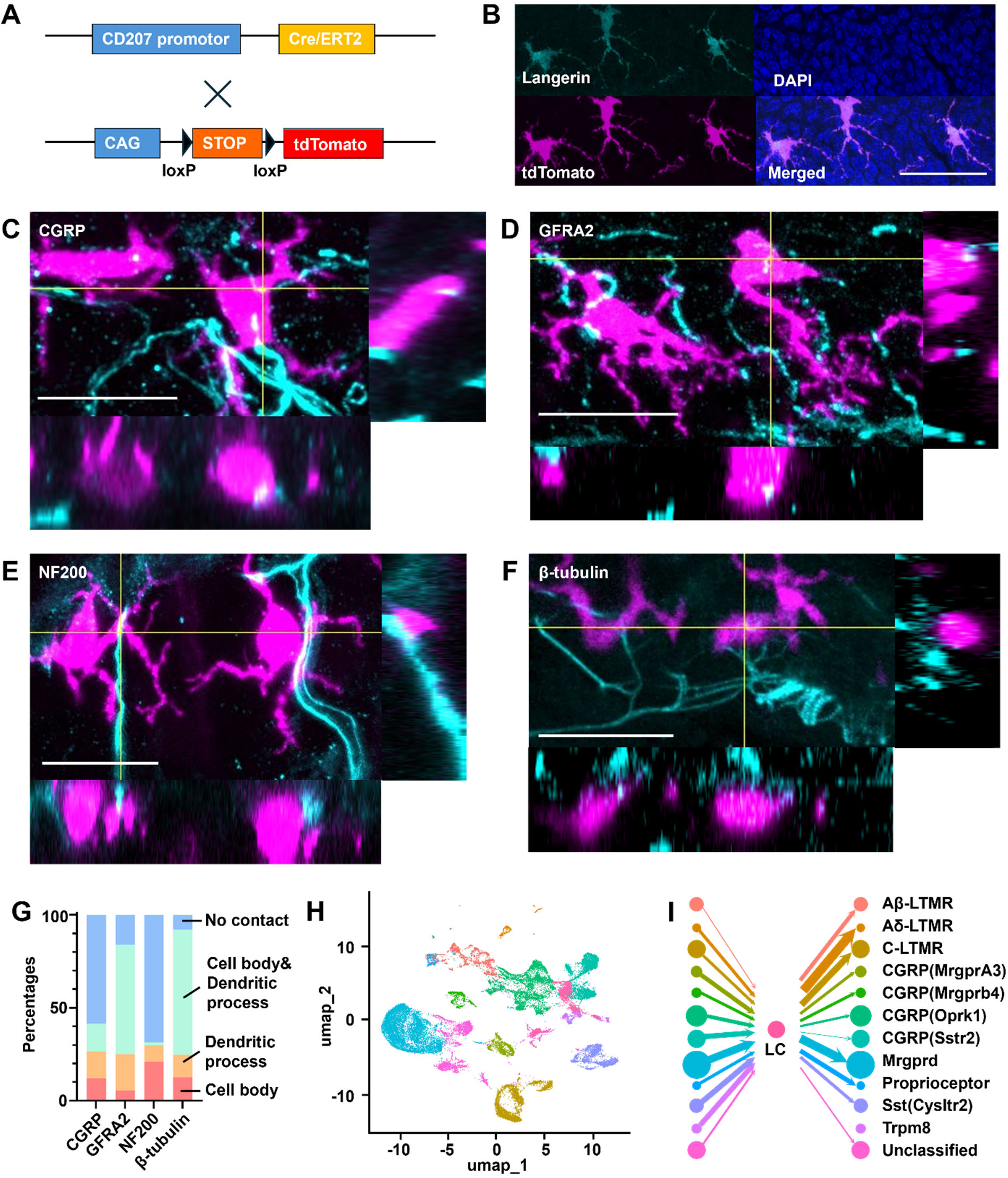
LCs preferentially communicate with non-peptidergic sensory neurons. **(A)** Genetic strategy for tamoxifen-inducible, LC-specific expression of tdTomato. In Lang^CreERT2^; Ai14 mice, Cre-ERT2 is expressed under control of the CD207 (Langerin) promoter. Tamoxifen administration induces nuclear Cre translocation and excision of a loxP-flanked STOP cassette, enabling tdTomato expression in LCs. **(B)** Representative immunofluorescence image of Lang^CreERT2^; Ai14 mouse skin showing strong overlap between tdTomato and Langerin, confirming selective labeling of LCs (*n* = 2-3 sections/mouse from 4 mice). Scale bar = 30 µm. **(C-G)** Spatial relationship between LCs and cutaneous sensory nerve fiber subtypes. Representative confocal images with orthogonal projections show proximity between tdTomato^+^ LCs (magenta) and nerve fibers (cyan) labeled for CGRP (peptidergic: C), GFRα2 (non-peptidergic: D), NF200 (myelinated A-fibers: E), and Pan-neuronal βIII-tubulin (F). Contact categories are defined as: LC cell body only (red), dendritic process only (orange), both cell body and dendritic process (green), or no contact (blue). Quantification shown in corresponding bar graphs (G: *n* = 3-5 sections/mouse from 5-6 mice). Scale bar = 20 µm. **(H)** UMAP visualization of integrated LC single-cell RNA-seq data with published DRG neuron transcriptomes. Cell populations are color-coded as in (I). **(I)** CellChat-inferred intercellular communication networks between LCs and DRG neuron subtypes. Node size corresponds to the relative cell population, while the thickness of arrow line indicates the strength of intercellular interactions. LCs exhibit the strongest bidirectional signaling with MrgprD-expressing non-peptidergic neurons.

To complement these anatomical findings with a transcriptomic interaction analysis, we applied CellChat, a mathematical modeling framework that infers intercellular communication networks based on ligand-receptor expression (*14*). Single-cell RNA sequencing data from LCs were integrated with published dorsal root ganglion (DRG) neuron transcriptomes (*15*), followed by clustering and visualization using uniform manifold approximation and projection (**Fig. 1H**). This combined dataset revealed at least 11 transcriptionally distinct sensory neuron clusters defined by established marker genes (*15*). CellChat analysis revealed differential interaction strengths between LCs and neuronal subtypes, visualized as weighted lines (**Fig. 1I**). The strongest incoming and outgoing interactions were observed between LCs and the MrgprD-expressing non-peptidergic neuronal cluster. Thus, anatomical and transcriptomic analyses independently identify non-peptidergic sensory neurons as a prominent LC-associated population, while also indicating that LCs reside within a broader network of cutaneous sensory afferents.

### LC activation potentiates sensory neuron responses without direct excitation

The anatomical association between LCs and sensory fibers raised the possibility that LC activity directly regulates sensory neuron function. To test this, we generated Lang^CreERT2^; Ai32 mice, in which LCs express a tamoxifen-inducible, improved channelrhodopsin-2-EYFP fusion protein (**Fig. 2A**). Following tamoxifen treatment, 94.8% (145/153) of LCs expressed ChR2-EYFP (CD207/ ChR2-EYFP = 97.3% (145/149); *n* = 2 sections/mouse from 4 mice; **Fig. 2B**). *In vivo* calcium imaging of DRG neurons revealed that blue light stimulation of the skin did not directly induce calcium influx in sensory neurons (0/230 cells from 3 mice; **Movie S1**). This lack of direct activation was further confirmed by electrophysiological recordings, which showed no increase in DRG firing frequency upon LC stimulation (**Fig. S3**). Next, we investigated whether LC activation modulates sensory neuron excitability. Blue light stimulation of LCs significantly increased both the proportion of heat-responsive DRG neurons and the magnitude of their responses to heat stimuli (**Fig. 2C-F**; **Movie S2**). A similar increase was observed in the proportion of neurons responding to mechanical stimulation, while the increase in the magnitude of mechanical responses was moderate yet statistically significant (**Fig. 2D,G,H; Movie S3**). Electrophysiological recordings further demonstrated that LC activation increased firing frequency in C-fibers, but not Aδ-fibers, during mechanical stimulation (**Fig. S3**). In control mice (Lang^Cre-negative^; Ai32 mice), blue light had no effect, indicating that the observed changes were dependent on LC-specific ChR2 expression and not due to nonspecific light effects. Additionally, LC activation enhanced neuronal responsiveness to chloroquine, a known agonist of MrgprA3-expressing pruriceptors, indicating increased itch-related signaling (**Fig. 2D**; **Movie S4**). Together, these findings suggest that while LC activation does not directly excite sensory neurons, it modulates their responsiveness to thermal, mechanical, and pruritic stimuli.

**Fig. 2.**
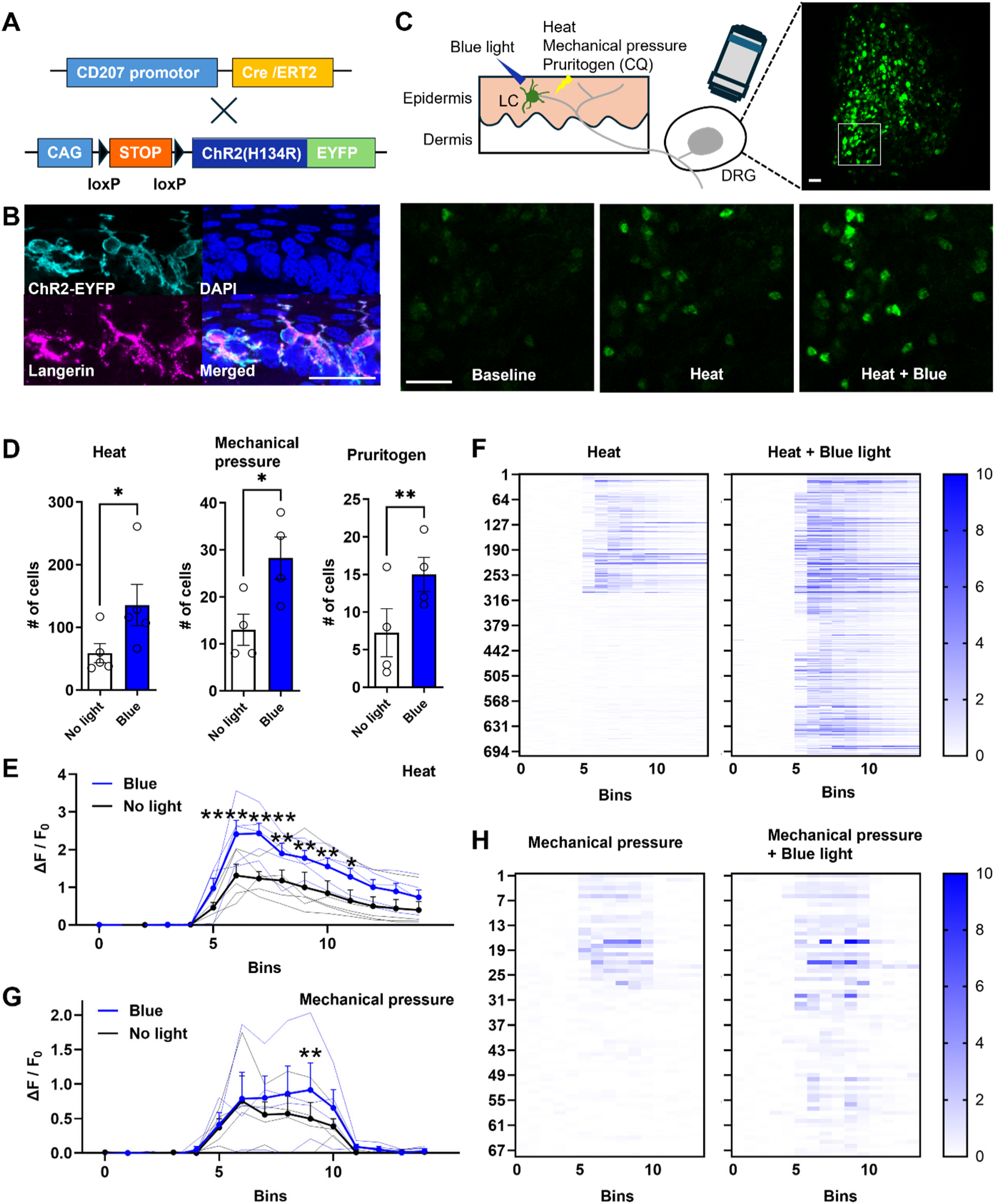
LC activation potentiates sensory neuron responses without direct excitation. **(A)** Genetic strategy for tamoxifen-inducible, LC-specific expression of channelrhodopsin-2 (ChR2-EYFP). In Lang^CreERT2^; Ai32 mice, tamoxifen-induced Cre recombination drives ChR2 expression selectively in LCs. **(B)** Immunofluorescence of mouse skin showing co-localization of ChR2-EYFP (cyan) and Langerin (magenta), confirming selective ChR2 expression in LCs (*n* = 2 sections/mouse from 4 mice). Scale bar = 30 µm. **(C)** Schematic (top left) illustrates the experimental design. LCs residing in the epidermis are in close proximity to sensory nerve terminals whose cell bodies are located in the DRG. Blue light activates ChR2-expressing LCs. Heat, mechanical pressure, or the pruritogen chloroquine (CQ) were applied with or without blue light. DRG neurons express GCaMP6s for calcium imaging. Representative fluorescence image of *in vivo* calcium imaging from DRG neurons (top right) with the boxed region expanded in the lower panels. Scale bar, 100 μm. Representative calcium imaging frames from the boxed region are shown at baseline (no stimulus), during heat application, and during combined heat and blue light stimulation (Heat + Blue), demonstrating enhanced neuronal calcium responses upon optogenetic co-activation (Movie S2). Scale bar, 100 μm. (D) Number of responding DRG neurons per preparation during blue light versus no-light conditions for heat (left), mechanical pressure (middle; Movie S3), and CQ (right; Movie S4) stimulation. Blue light significantly increased the number of responding neurons under all three modalities. Data shown as mean ± SEM; circles represent individual animals. \**p* < 0.05, \*\**p* < 0.01 (heat and mechanical pressure: paired, two-tailed Student’s *t* test, *n*=4-5 mice; CQ: unpaired two-tailed Student’s *t* test, *n*=4 mice). (E) Mean ΔF/F₀ traces (± SEM) for heat responding neurons during heat stimulation with (blue) or without (black) concurrent blue light. Stimulation was delivered during bins 5-8. Thin lines represent individual animal averages. Each bin indicates 7.5-8.5 seconds. Asterisks denote time bins with significant differences between conditions (\**p* < 0.05, ** *p* < 0.01, **** *p* < 0.0001). two-way repeated-measures ANOVA with Bonferroni’s post hoc test, F (14,56)=4.968, *p*<0.0001. *n*=5 mice. (F) Heatmaps displaying ΔF/F₀ responses of individual DRG neurons (rows) across imaging time bins (columns) during heat alone (left) and heat + blue light (right). Each row represents one neuron; color scale indicates ΔF/F₀ magnitude (0–10). LC optogentic stimulation recruits a larger population of neurons and increases response amplitude. (G) Mean ΔF/F₀ traces during mechanical pressure stimulation with (blue) or without (black) blue light, plotted as in (E). \*\**p* < 0.01. two-way repeated-measures ANOVA with Bonferroni’s post hoc test, F (14,42)=2.052, *p*=0.0366. *n*=4 mice. (H) Heatmaps as in (F) for mechanical pressure alone (left) and mechanical pressure + blue light (right).

### LC activation enhances nociceptive and pruritic behaviors

We next asked whether LC-dependent enhancement of sensory neuron responsiveness translates into altered behavior. To selectively activate LCs *in vivo*, we generated Lang^CreERT2^; Gq-coupled DREADD (designer receptors exclusively activated by designer drugs) mice (**Fig. 3A**). Following tamoxifen treatment, 97.1% (365/376) of LCs expressed HA-tagged Gq DREADD (CD207/HA = 99.2% (365/368); *n* = 2 sections/mouse from 3 mice; **Fig. 3B**). While PBS did not alter thermal withdrawal latencies in either Lang^CreERT2^; Gq-coupled DREADD mice or control mice (Lang^Cre-negative^; Gq-coupled DREADD mice; **Fig. 3C**), the DREADD ligand JHU37160 (*16*) decreased paw withdrawal latency in Lang^CreERT2^; Gq-coupled DREADD mice (**Fig. 3D**), indicating LC activation increases heat sensitivity. Similarly, PBS did not alter mechanical paw withdrawal threshold in either Lang^CreERT2^; Gq-coupled DREADD mice or control mice (**Fig. 3E**), the DREADD ligand decreased paw withdrawal threshold in Lang^CreERT2^; Gq-coupled DREADD mice (**Fig. 3F**), suggesting LC activation increases mechanical sensitivity. LC activation alone did not induce spontaneous scratching (**Fig. 3G**), indicating that LC activity is not sufficient to generate itch behavior in the absence of a pruritic stimulus. However, LC activation significantly increased scratching evoked by histamine, chloroquine, and serotonin (**Fig. 3H–J**). There are no significant sex differences in thermal and mechanical withdrawal thresholds and scratching (**Fig. S4**).

**Fig. 3.**
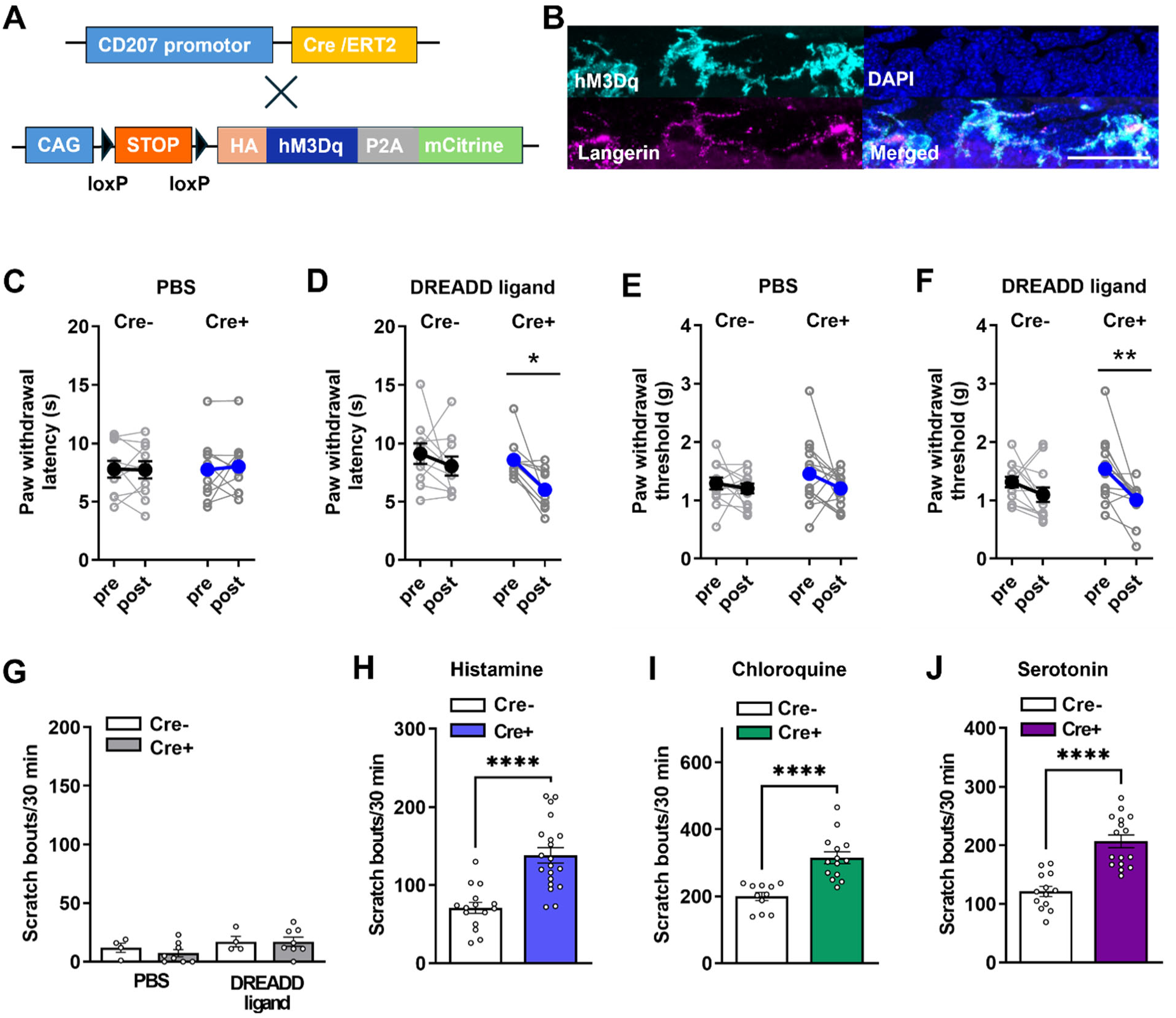
Chemogenetic activation of LCs enhances nociception and itch. **(A)** Genetic strategy for tamoxifen-inducible, LC-specific expression of hM3Dq-HA (Gq-DREADD). In Lang^CreERT2^; Gq-DREADD mice, tamoxifen-induced Cre recombination drives Gq-DREADD expression selectively in LCs. **(B)** Immunostaining for HA tag (cyan) and Langerin/CD207 (magenta) confirming Gq-DREADD expression in LCs (*n* = 2 sections/mouse from 3 mice). Scale bar = 30 µm. **(C-D)** Paw withdrawal latency to radiant heat before (pre) and after (post) injection of PBS (C) or the DREADD agonist JHU37160 (D) in Cre⁻ and Cre⁺ mice. JHU37160, but not PBS, significantly reduces withdrawal latency in Cre⁺ mice. *n* = 13-14 per group. Two-way repeated-measures ANOVA with Bonferroni’s post hoc test: PBS, F(1,25)=2.266, *p*=0.1448; JHU37160, F(1,25)=15.11, *p*=0.0007. **(E-F)** Mechanical paw withdrawal threshold before and after injection of PBS (E) or JHU37160 (F). JHU37160 selectively reduces mechanical threshold in Cre⁺ mice; PBS has no effect. *n* = 10 per group. Two-way repeated-measures ANOVA with Bonferroni’s post hoc test: PBS, F(1,18)=0.000, *p*=0.8231; JHU37160, F(1,18)=7.497, *p*=0.0135. **(G)** LC activation alone does not induce spontaneous scratching. *n* = 4-8; two-way repeated-measures ANOVA with Bonferroni’s post hoc test, F(1,10)=0.2925, *p*=0.6005 **(H-J)** LC activation enhances scratching behavior evoked by intradermal injection of histamine (H), chloroquine (I), and serotonin (J) in Cre⁺ versus Cre⁻ mice. *n* = 15-20 (histamine), 11-14 (chloroquine), 13-16 (serotonin); unpaired two-tailed Student’s *t* test; all comparisons *p* < 0.0001. \**p* < 0.05, \*\**p* < 0.01, \*\*\*\**p* < 0.0001.

To further test whether activation of LCs could enhance nociception and pruriception, we used Lang^CreERT2^; Ai32 mice. While either blue or green light did not alter thermal withdrawal latency or mechanical withdrawal threshold in control mice (tamoxifen-non treated mice; **Fig. S5A, S5C**), the blue light, but not green light, decreased paw withdrawal latency and mechanical withdrawal threshold in tamoxifen-treated mice (**Fig. S5B, S5D**). Blue light did not elicit scratching, but enhanced scratching behavior evoked by histamine and chloroquine (**Fig. S5E-G**). Thus, two independent approaches to LC activation demonstrate that LC activity amplifies stimulus-evoked nociceptive and pruritic behaviors without itself generating spontaneous itch behavior.

### LC activity is required for pruriception but not baseline nociception

The activation experiments established that LC activity is sufficient to potentiate sensory responses. We next asked whether endogenous LC activity is required for pain and itch behavior. We generated Lang^CreERT2^; Gi-coupled DREADD mice, in which LCs could be selectively inhibited by chemogenetic stimulation (**Fig. 4A**). Following tamoxifen treatment, 98.4% (240/244) of LCs expressed HA-tagged Gi DREADD (CD207/HA = 98.8% (240/243); *n* = 2-3 sections/mouse from 3 mice; **Fig. 4B**). Neither PBS nor the DREADD ligand altered thermal withdrawal latencies in either Lang^CreERT2^; Gi-coupled DREADD mice or control mice (Lang^Cre-negative^; Gi-coupled DREADD mice) (**Fig. 4C-D**). Similarly, PBS or the DREADD ligand did not alter mechanical paw withdrawal threshold in either Lang^CreERT2^; Gi-coupled DREADD mice or control mice (**Fig. 4E-F**). These results suggest that LCs are not required for basal heat and mechanical sensitivities, consistent with a previous finding showing that LCs depleted mice exhibited unchanged basal heat and mechanical sensitivities (*10*). In contrast, the DREADD ligand significantly reduced scratching evoked by histamine, chloroquine and serotonin (**Fig. 4G-I**).

**Figure 4.**
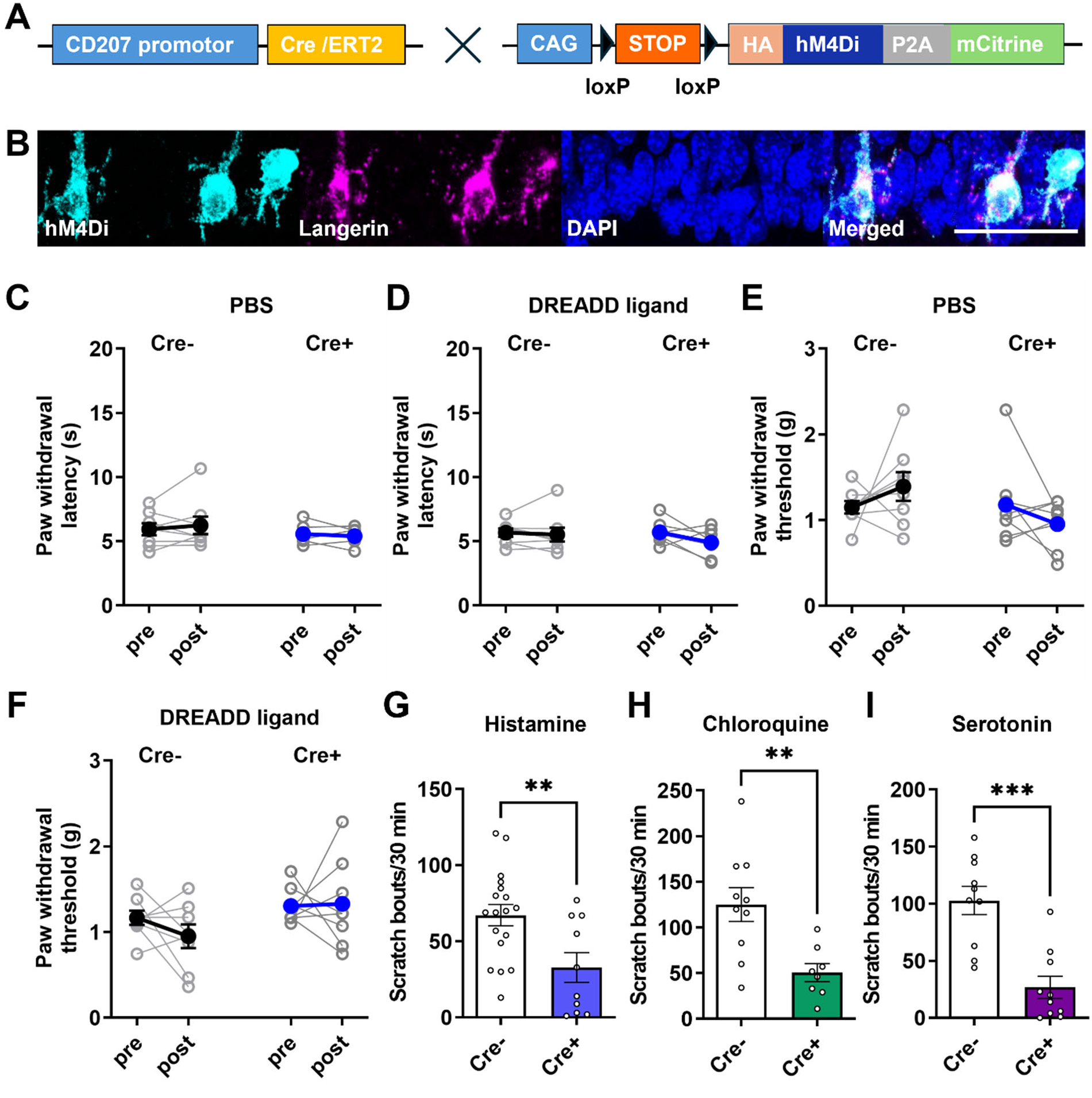
LCs are required for pruritic but not basal thermal or mechanical responses. **(A)** Genetic strategy for tamoxifen-inducible, LC-specific expression of hM4Di-HA (Gi-DREADD). In Lang^CreERT2^; Gi-DREADD mice, tamoxifen-induced Cre recombination drives Gi-DREADD expression selectively in LCs. **(B)** Immunostaining for HA tag (cyan) and Langerin/CD207 (magenta) confirming Gi-DREADD expression in LCs (*n* = 2-3 sections/mouse from 3 mice). Scale bar, 30 µm. **(C-D)** Paw withdrawal latency to radiant heat before (pre) and after (post) injection of PBS (C) or JHU37160 (D) in Cre⁻ and Cre⁺ mice. Chemogenetic LC inhibition does not alter thermal withdrawal latency. *n* = 7-8 per group. Two-way repeated-measures ANOVA with Bonferroni’s post hoc test; PBS, F (1,13)=0.04514, *p*=0.835; JHU37160, F (1,13)=2.451, *p*=0.1414. **(E-F)** Mechanical paw withdrawal threshold before and after injection of PBS (E) or JHU37160 (F). LC inhibition does not affect baseline mechanical sensitivity. *n* = 7-8 per group. Two-way repeated-measures ANOVA with Bonferroni’s post hoc test; PBS, F (1,14)=0.004504, *p*=0.9474; JHU37160, F (1,14)=0.4589, *p*=0.5092. **(G-I)** Chemogenetic LC inhibition significantly reduces scratching evoked by intradermal injection of histamine (G), chloroquine (H), or serotonin (I) in Cre⁺ versus Cre⁻ mice. *n* = 10-18 (histamine), 8-10 (chloroquine), 10 (serotonin); unpaired two-tailed Student’s *t* test: Histamine, *p*=0.0073; chloroquine, *p*=0.0045; serotonin, *p*=0.0001. \*\**p* < 0.01. \*\*\**p* < 0.001.

To further confirm this finding, we used Lang-DTR mice, in which LCs can be selectively ablated by diphtheria toxin. DTR-tagged EGFP was expressed in 72.1% of LCs (194/269), with high specificity for CD207⁺ cells (CD207⁺/DTR⁺ = 96.5%; *n* = 2 sections/mouse from 3 mice; **Fig. S6B**). Diphtheria toxin administration resulted in efficient LC depletion that persisted for at least 7 days (**Fig. S6A, S6C, and S6D**), consistent with previous reports (*10, 17*). Lang-DTR mice receiving diphtheria toxin exhibited significantly reduced scratching responses to histamine, serotonin, and chloroquine (**Fig. S6E-G**), indicating that LCs are required for pruritic behavior.

We next asked whether LCs can directly detect pruritogenic signals. *In vitro* calcium imaging of cultured LCs revealed responses to histamine, serotonin, and chloroquine (**Fig. S7A-B**). Of 406 LCs examined, 10% (41/406) responded to at least one pruritogen. Response profiles were largely non-overlapping (**Fig. S7C**). These findings demonstrate that a subset of LCs can directly respond to distinct pruritogens, providing a potential cellular mechanism through which local immune sensing could influence nearby sensory neurons.

### LCs express semaphorin7A, which potentiates pruriceptive signaling

We next sought to identify molecular signals that could mediate communication between LCs and sensory neurons. CellChat analysis identified several candidate ligand-receptor interactions, including Cxcl2, Wnt4, Cdh1, Nectin1, Sema7a, and Thbs1 (**Table S1**). Because these predictions are based on transcriptomic expression and are therefore hypothesis-generating, we selected Sema7a for functional validation. Immunostaining of mouse skin from Lang^CreERT2^; Ai14 mice showed that 24.0% of LCs (196/818; *n*= 3 slides/mouse from 3 mice) expressed semaphorin7A (**Fig. 5A**). Staining of isolated LCs gave a similar result, with 26.4% of cells semaphorin7A^+^ (37/140; *n* = 2 slides/mouse from 3 mice; **Fig. 5B**). These data identify a subset of LCs that could provide semaphorin7A-dependent signals to neighboring sensory fibers.

**Fig. 5.**
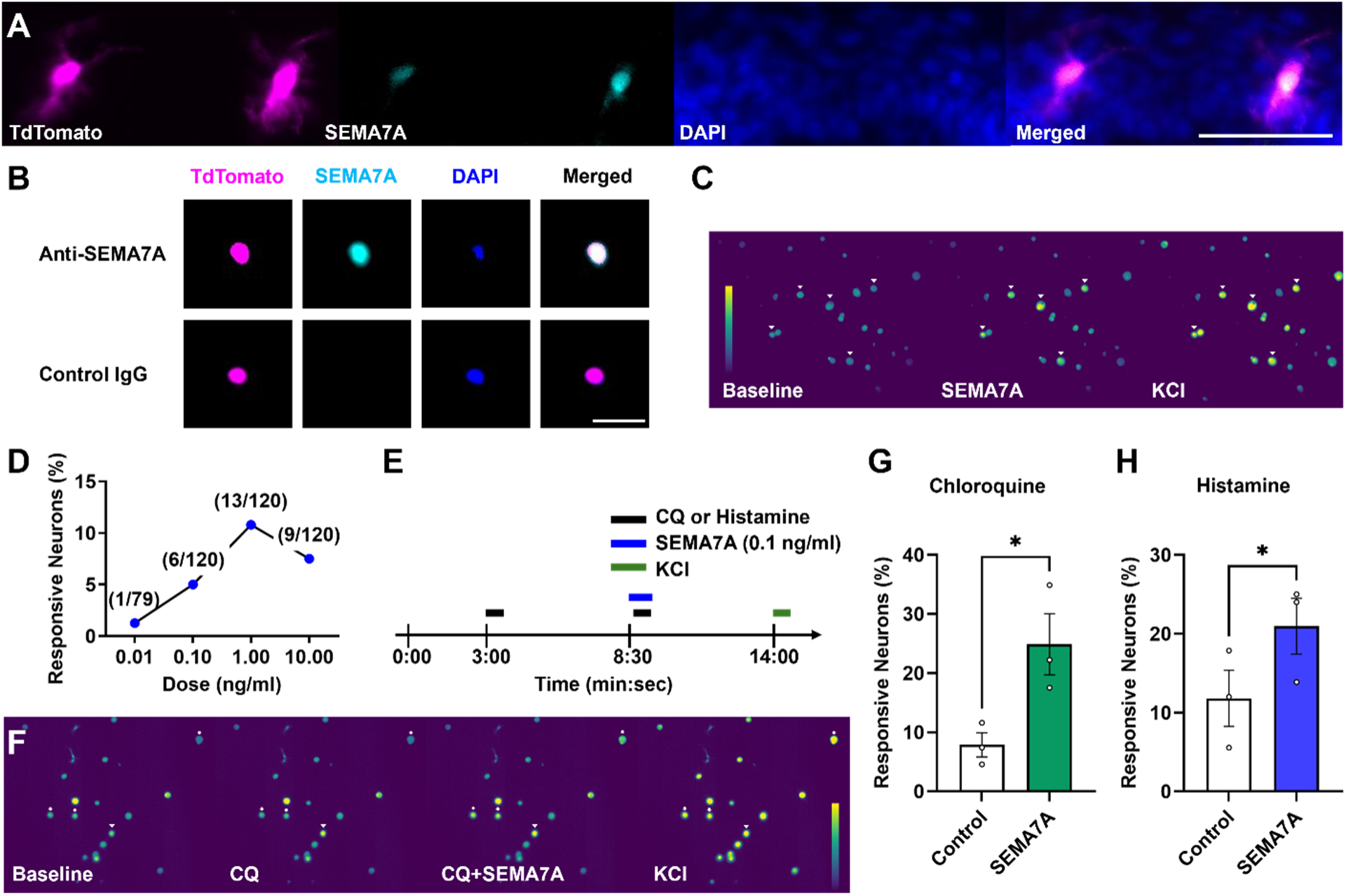
LCs express semaphorin7A, which sensitizes pruriceptive sensory neurons. (A) Representative immunofluorescence image of skin sections from Lang^CreERT2^; Ai14 mouse, in which LCs express tdTomato (magenta), stained for semaphorin 7A (SEMA7A; cyan) and DAPI (*n* = 3 sections/mouse from 3 mice). Scale bar = 50 µm. (B) Representative images of cultured LCs from Lang^CreERT2^; Ai14 mice expressing TdTomato (magenta), stained for SEMA7A (cyan), and DAPI (blue), probed with anti-SEMA7A antibody or control IgG. Scale bar, 50 μm. *n* = 2-3 slides/mouse from 3 mice. (C) Representative pseudocolor calcium imaging traces of dissociated DRG neurons at baseline, following SEMA7A application, and following KCl application. Arrowheads indicate SEMA7A -responsive neurons. (D) Dose-response curve of the percentage of DRG neurons responsive to increasing concentrations of SEMA7A (0.01-10 ng/ml). Numbers above each point indicate responsive neurons/total neurons tested at that dose. *n* = 2-3 dishes/mouse from 3 mice. (E) Schematic of the stimulation protocol for calcium imaging in (F) to (H): chloroquine (CQ) or histamine, SEMA7A (0.1 ng/ml), and KCl were applied at the indicated times. (F) Representative pseudocolor calcium imaging traces of DRG neurons at baseline and following sequential application of CQ, CQ + SEMA7A, and KCl. Arrowheads indicate CQ-responsive neurons. Arrows indicate neurons responding only to CQ + SEMA7A (not to CQ alone). (G) Percentage of DRG neurons responsive to chloroquine alone (Control) or to chloroquine plus SEMA7A. Neurons were first exposed to CQ, followed by co-application of CQ and SEMA7A. (H) Percentage of DRG neurons responding to histamine alone (Control) or to histamine plus SEMA7A, using the same sequential application protocol. In (G) and (H), each point represents 2-3 dishes/mouse from one animal; *n* = 3 mice. Data shown as mean ± SEM. (G) *p*=0.033; (H) *p*=0.0245; paired two-tailed Student’s *t* test.

To determine whether semaphorin 7A directly modulates sensory neuron activity, we performed calcium imaging in primary-cultured DRG neurons. Bath application of semaphorin 7A elicited responses in a concentration-dependent, bell-shaped manner across the tested concentrations (0.01-10 ng/ml), with the highest proportion of responsive neurons observed at 1.0 ng/ml (13/120, 10.8%) (**Fig. 5C-D**). We next asked whether semaphorin 7A enhances neuronal responsiveness to chloroquine or histamine. To minimize potential ceiling effects, we used relatively low concentrations of semaphorin 7A (0.1 ng/ml), chloroquine (100 μM) and histamine (3 μM). Following an initial application of chloroquine or histamine, semaphorin 7A was applied before a second challenge with the same agonist (**Fig. 5E**). Repeated application of chloroquine or histamine alone did not significantly alter the proportion of responsive neurons (chloroquine: 8.5% for the first application versus 10.9% for the second; histamine: 13.2% versus 12.2%). In contrast, pretreatment with semaphorin 7A significantly increased the proportion of neurons responding to both chloroquine and histamine (chloroquine: 7.9% versus 24.9%; histamine: 11.8% versus 21.0%) (**Fig. 5F-H**). These findings indicate that semaphorin 7A enhances sensory neuron responsiveness to both chloroquine and histamine.

### Pruriceptive sensory neurons express PlexinC1, which mediates semaphorin7A-induced responses

We next examined expression of the semaphorin7a receptor PlexinC1 in DRG neurons. Immunostaining of DRG sections revealed robust PlexinC1 expression, including within a subset of neurons expressing the non-peptidergic marker IB4 (**Fig. 6A**) and MrgprA3, a marker of pruriceptive sensory neurons (**Fig. 6B**). Quantification showed that PlexinC1 substantially overlapped with IB4 (78.1% (750/960); *n* = 2-3 sections/mouse from 3 mice; **Fig. 6C**) and, notably, with MrgprA3⁺ pruriceptors, the majority of which co-expressed PlexinC1 (83.4% (685/821); *n* = 2-3 sections/mouse from 3 mice; **Fig. 6D**). These findings indicate that PlexinC1 is broadly expressed across itch-sensing subpopulations of DRG neurons, anatomically positioning these neurons to respond to semaphorin 7A signals in the local microenvironment.

**Fig. 6.**
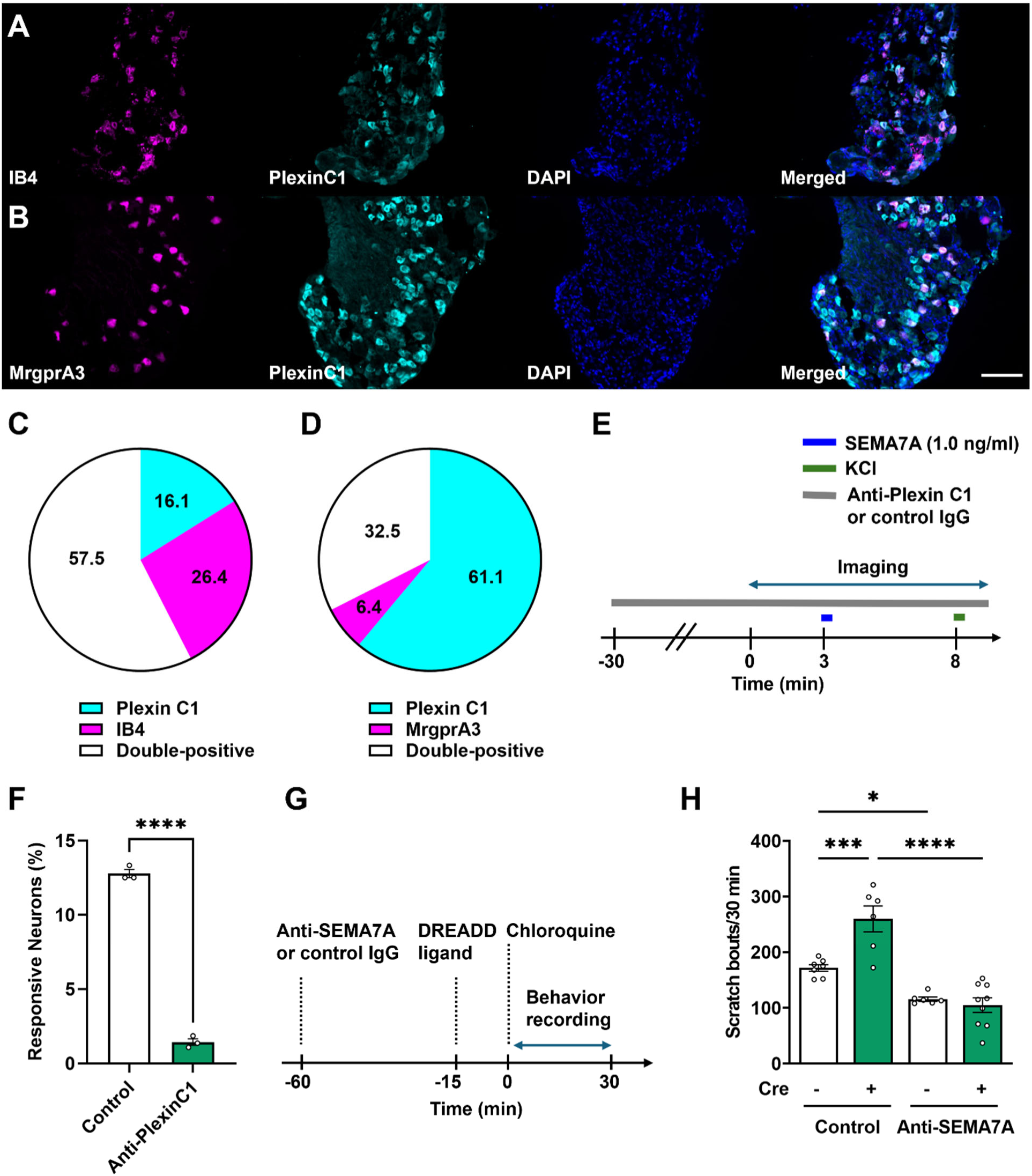
Pruriceptive sensory neurons express PlexinC1, which mediates semaphorin7A-induced responses. (A) Representative images of DRG sections stained for IB4 (magenta), PlexinC1 (cyan), and DAPI (blue). (B) Representative images of DRG sections from MrgprA3^Cre^; Ai14 mouse, in which MrgprA3^+^ neurons express tdTomato (magenta), stained for PlexinC1 (cyan), and DAPI (blue). Scale bar, 100 μm (applies to A and B). (C) Quantification of overlap between PlexinC1⁺ and IB4⁺ DRG neurons, expressed as percentage of all neurons positive for either marker; *n* = 2-3 sections/mouse from 3 mice. (D) Quantification of overlap between PlexinC1⁺ and MrgprA3⁺ DRG neurons, as in (C). (E) Schematic of *in vitro* calcium imaging protocol. DRG neurons were incubated with anti-PlexinC1 antibody or control IgG starting at -30 min, and the antibody was present throughout imaging. Semaphorin 7A (SEMA7A; 1.0 ng/ml) was applied at 3 min and KCl at 8 min. (F) Percentage of neurons responding to SEMA7A in the presence of control IgG or anti-PlexinC1. Each point represents 2-3 dishes/mouse from one animal; *n* = 3 mice. Data shown as mean ± SEM. \*\*\*\**p* < 0.0001; unpaired two-tailed Student’s *t* test. (G) Schematic of the *in vivo* behavioral protocol: anti-SEMA7A or control IgG was administered at -60 min, DREADD ligand, JHU37160, at -15 min, and chloroquine at 0 min, followed by 30 min of scratching behavior recording in Lang^CreERT2^; Gq-DREADD mice and Cre-negative littermates. (H) Number of scratch bouts over 30 min following chloroquine challenge in Cre⁻ and Cre⁺ mice pretreated with control IgG or anti-SEMA7A. *n* = 6-9 per group. Data shown as mean ± SEM. \**p* < 0.05, \*\*\**p* < 0.001, \*\*\*\**p* < 0.0001; two-way ANOVA with Tukey’s post hoc test. F (1,24)=13.04, *p*=0.0014.

To test whether PlexinC1 mediates the neuronal response to semaphorinA7A, we performed calcium imaging of cultured DRG neurons in the presence of an anti-PlexinC1 antibody or control IgG (**Fig. 6E**). We used semaphorin7A at 1.0 ng/ml, the concentration that evoked the largest response (**Fig. 5D**), so that a reduction in responsiveness could be detected. Anti-PlexinC1 significantly reduced the proportion of semaphorin7A-responsive neurons (12.8% vs 1.4%; 2-3 dishes/mouse from 3 mice; unpaired *t-test*, p<0.0001; **Fig. 6F**).

Finally, we tested whether semaphorin 7A is required for LC-dependent potentiation of itch *in vivo*. We generated Lang^CreERT2^; Gq-coupled DREADD mice to enable chemogenetic activation of LCs and administered a Sema7A-neutralizing antibody or control IgG prior to DREADD ligand and chloroquine challenge (**Fig. 6G**). In control IgG-treated animals, chemogenetic activation of LCs (Cre^+^) significantly increased chloroquine-evoked scratching compared with Cre-negative littermates (**Fig. 6H**). This increase was abolished by anti-Sema7A. Cre^+^ mice given anti-Sema7A scratched at levels similar to Cre^-^mice given anti-Sema7A, and significantly less than Cre^+^ mice given control IgG (**Fig. 6H**). Anti-Sema7A also reduced scratching in Cre⁻ mice, which suggests that semaphorin 7A contributes to chloroquine-evoked itch even without LC activation. Together, these results indicate that semaphorin 7A is required for the enhancement of chloroquine-evoked itch caused by LC activation, consistent with semaphorin 7A acting on PlexinC1-expressing sensory neurons.

## DISCUSSION

Our findings identify epidermal LCs as regulators of cutaneous sensory gain. LCs lie within a dense network of sensory fibers and are preferentially associated with non-peptidergic afferents. Activating them increased neuronal and behavioral responses to thermal, mechanical, and pruritic stimulus, but did not excite sensory neurons on their own. Inhibiting or removing LCs reduced pruritogen-evoked scratching and left baseline thermal and mechanical sensitivity unchanged. LCs therefore behave as modulators of sensory responsiveness rather than primary sensory transducers, and they are specifically required for normal itch. We also identified a molecular pathway for this modulation. Semaphorin7A, expressed by a subset of LCs, sensitizes pruriceptive neurons that express PlexinC1, and blocking semaphorin7A abolished the LC-driven enhancement of chloroquine-evoked itch.

The anatomical and transcriptomic data converge on preferential, but not exclusive, association between LCs and non-peptidergic sensory neurons. LC cell bodies and dendrites were more frequently associated with GFRα2^+^ non-peptidergic fibers than with CGRP^+^ peptidergic or NF200^+^ myelinated fibers, while CellChat analysis identified the MrgprD^+^ non-peptidergic cluster as having the strongest predicted interactions with LCs. However, a substantial fraction of LC processes remained associated with peptidergic fibers, indicating that LCs are positioned to interact with multiple sensory neuron populations rather than a single dedicated neuronal subtype. Two considerations are important when interpreting these anatomical findings. First, GFRα2 and MrgprD identify overlapping but non-identical neuronal populations. Thus, our results support preferential association with the broader non-peptidergic class rather than a single genetically defined neuronal subtype. Second, our confocal analyses establish spatial proximity between LCs and axonal profiles but do not demonstrate direct membrane contact or a synaptic-like interaction. These relationships should therefore be interpreted within the spatial resolution of the imaging approach.

This anatomical partial selectivity is reflected in the broad functional effects we observed. Optogenetic and chemogenetic activation of LCs enhanced neuronal and behavioral responses not only to chloroquine (an MrgprA3-dependent, non-histaminergic pruritogen) but also to histamine and serotonin, and additionally sensitized neurons to thermal and mechanical stimuli. Conversely, chemogenetic inhibition or diphtheria toxin-mediated ablation of LCs attenuated scratching evoked by all three pruritogens tested. This contrasts with the more restricted phenotype reported after selective ablation of MrgprD⁺ neurons, which reduces β-alanine-evoked itch without affecting chloroquine responses (*18*), and instead more closely resembles the broader deficits seen following ablation of MrgprA3^+^ neurons (*19*) or peptidergic afferents (*20*). We note that our functional readouts, *in vivo* calcium imaging, electrophysiology, and behavioral assays, were performed in heterogeneous DRG neuron populations or at the whole-animal level, without genetic or molecular resolution of neuronal subtype. Our data therefore more directly support a model in which LC activation broadly increases sensory neurons responsiveness and pruriceptive and nociceptive output rather than one in which LC signaling is confined to a single neuronal lineage.

Two observations from our functional experiments help explain how a single immune cell type could exert such broad effects. First, optogenetic stimulation of LCs did not itself evoke calcium responses or increase firing in sensory neurons, indicating that LCs are not acting as primary transducers of pruritic or nociceptive stimuli. Second, *in vitro* calcium imaging showed that a subset of LCs responds directly to histamine, serotonin, and chloroquine, with largely non-overlapping response profiles across agonists. Together, these findings support a model in which LCs can detect local pruritogenic signals and, through subsequent interactions with nearby sensory afferents, amplify sensory responses within the cutaneous microenvironment. Thus, rather than generating sensory activity directly, LCs may function as local immune-derived modulators that adjust the responsiveness of cutaneous sensory circuits to incoming stimuli.

The physiological relevance of this LC-dependent modulation is supported by complementary loss-of-function approaches. Both acute chemogenetic inhibition of LC activity using Gi-DREADD and genetic depletion using Lang-DTR significantly reduced scratching evoked by histamine, chloroquine, and serotonin, while leaving basal thermal and mechanical sensitivity unchanged. Depleting cells with diphtheria toxin can alter epidermal homeostasis or local immune composition, but the concordance with acute, reversible chemogenetic inhibition argues that the phenotype reflects loss of LC function rather than tissue disruption. The contrast between activation and loss of function is informative. Activating LCs raised pain-related sensitivity, yet removing them did not lower baseline pain sensitivity, a result consistent with earlier work (*10*). LCs thus appear to modulate itch under basal conditions, while their effect on pain may emerge only when they are activated, as might occur in injury or inflammation.

Our CellChat analysis nominated several candidate mediators of this LC-to-neuron signaling axis (Table S1). Among these, we prioritized semaphorin 7A for full experimental validation and identified it as a functionally necessary mediator of LC-dependent itch potentiation. Semaphorin 7A was expressed in roughly a quarter of LCs, directly activated cultured DRG neurons in a concentration-dependent manner, and sensitized neuronal responses to both chloroquine and histamine at subthreshold concentrations. Its receptor, PlexinC1, was broadly expressed across non-peptidergic (IB4⁺) and pruriceptive (MrgprA3⁺) DRG neuron populations, and an anti-PlexinC1 antibody sharply reduced semaphorin7A-evoked responses. Neutralizing semaphorin 7A *in vivo* eliminated the chemogenetic LC activation-induced enhancement of chloroquine-evoked scratching, demonstrating that semaphorin 7A is required for LC-mediated itch potentiation, consistent with signaling via its receptor PlexinC1.

The physiological and pathological significance of this LC-dependent sensory modulation remains to be determined. Because LC density and activation state are dynamically altered in inflammatory skin diseases such as atopic dermatitis (*21–24*) and psoriasis (*25–27*), as well as painful peripheral neuropathy, such as chemotherapy-induced neuropathy (*28, 29*) and diabetic-induced neuropathy (*30–32*), the LC-Semaphorin 7A-PlexinC1 axis we describe here may be relevant to disease-associated sensory dysfunction. Pathological expansion or activation of LCs could amplify cutaneous pain and itch signaling beyond what is driven by neuronal sensitization alone, raising the possibility that targeting LC-derived mediators, semaphorin 7A in particular, could offer a therapeutic strategy for chronic pruritus that is distinct from and complementary to neuron-targeted approaches. Whether LCs contribute to itch in these specific disease contexts, and whether the Semaphorin7A-PlexinC1 axis identified here is involved, remains an important direction for future investigation.

In summary, our results show that epidermal LCs adjust the responsiveness of cutaneous sensory circuits and are required for normal pruritogen-evoked itch. They also identify semaphorin7A as a mediator of LC-induced itch potentiation, with PlexinC1 on pruriceptive neurons as a candidate receptor.

## MATERIALS AND METHODS

### Study Design

This study used a controlled laboratory experimental design to investigate how LCs regulate sensory processing. We used multiple genetic mouse models, primary cultures of DRG neurons and LCs, and *in vivo* and *in vitro* experimental approaches to examine interactions between LCs and sensory neurons. Both male and female mice were used throughout the study, and age-matched littermates were used as controls whenever possible. Power analyses (α = 0.05, power = 0.8) were used to estimate target sample sizes, although final group sizes varied according to breeding success and litter size. Animal numbers and sample replicates are reported in the corresponding figure legends or Results section. Animals were randomly assigned to treatment groups. Investigators were blinded to treatment group during data collection, and observers were blinded to treatment group during video and image analysis. No data were excluded as outliers.

### Mice

All mouse procedures were approved by Institutional Animal Care and Use Committees of the University of Miami (protocol number: 21-111and TR202400000053). Lang^CreERT2^ (strain#:016940), Nav1.8^cre^ (strain#036564), Ai14 (strain#:007914), Ai32 (strain#:024109), Gq-coupled DREADD (strain#:026220), Gi-coupled DREADD (strain#:026219), Lang^DTR-EGFP^ (strain#:016940) and C57BL/6J mice were obtained from the Jackson Laboratory (Bar Harbor, ME). MrgprA3^Cre^ mice were provided by Dr. Xinzhong Dong, Johns Hopkins University (*19*). They were maintained as mating colonies within our facility. Mice were group housed (2-5 per cage) in a 12-12 light-dark cycle and had access to food and water *ad libitum*. The sample size was determined by power analysis. Adult male and female mice were randomly assigned to experimental groups. The experiment was replicated at least two times in laboratory. At the conclusion of experiments, mice were euthanized via transcardial perfusion or cervical dislocation under anesthesia.

### Scratch test

Prior to testing, mice were habituated for 5 days to the behavior recording chamber, handling, and intradermal (i.d.) injections (*33*). A small patch of fur on the nape of the neck was shaved for i.d. injections in all mice. Mice were administered i.d. injections of 10 μL of histamine (50 μg, Sigma-Aldrich, St. Louis, MO), chloroquine (100 μg, Sigma-Aldrich), or serotonin (10 μg, Sigma-Aldrich) and then recorded for 30 minutes. For optogenetic experiments, the light was delivered from the center of the ceiling of the recording arena (*34*). Behavior was recorded for 30 min with green light, blue light, or no light stimulation of the shaved skin (power: 2.5-7 mW, frequency: 20 Hz, distance between light and skin surface: 1-5 cm). For chemogenetic experiments, Lang^CreERT2^; Gq-coupled DREADD or Lang^CreERT2^; Gi-coupled DREADD mice were administered i.d. injections of 10 μL of JHU37160 dihydrochloride (25 μM, Hello Bio Inc., Princeton, NJ) 15 min before pruritogen injection (*16*). For neutralizing antibody experiments, Lang^CreERT2^; Gq-coupled DREADD mice were administered i.d. injections of 25 μL of anti-Sema7a antibody (200 μg/ml; AF1835; R&D Systems, Minneapolis, MN; RRID:AB_2186134) or control goat IgG (200 μg/ml; AB-108-C; R&D Systems; RRID:AB_354267) 60 min before pruritogen injection (*35, 36*). For conditional LC ablation experiments, Lang^DTR-EGFP^ mice were administered intraperitoneal injections of DT (1 μg in 100 μl, Sigma Aldrich) (*10*). Prior to each testing session, the experimental chamber was sanitized with 70% ethanol, both at the beginning of the day and after completion of testing for each group of mice to minimize odor effects. The number of scratch bouts was analyzed in 5-min bins by a trained observer blinded to the treatment condition. One scratch bout was defined as one or more rapid back-and-forth hind paw motions directed toward and contacting the injection site, ending with licking or biting of the toes or placement of the hind paw on the floor.

### Von Frey test

Mice were acclimated to a perforated metal floor for 120 min pre two days and 1 hour before testing (*37*). The plantar surface of the hind paws was tested with a series of von Frey filaments to determine the paw withdrawal threshold (PWT) by trained experimenters blinded to the treatment condition. 0.6 g (No. 3.84) filament was used for first application and von Frey filament intensity was lowered or increased for up-down test. The filament was kept contacting 3 seconds to the skin area after the filament bent. The 50% paw withdrawal threshold was calculated using the formula developed by Chaplan et al., (1994). For optogenetic experiments, the plantar surface of the hindpaws was tested with a series of von Frey filaments along with green light, blue light, or no light stimulation (4 mW, 20 Hz) (*34*). For chemogenetic experiments, Lang^CreERT2^; Gq-coupled DREADD or Lang^CreERT2^; Gi-coupled DREADD mice were administered i.d. injections of 5 μL of JHU37160 dihydrochloride (25 μM, Hello Bio Inc., Princeton, NJ) 15 min before testing (*16*).

### Hargreaves test

Mice were placed into plastic cage on heat glass at 30°C for habituation (*37*). Habituation was performed 2 hours continuous pre 2 days and 1 hour before thermal test. To determine the paw withdrawal latencies (PWLs), radiant beam was applied to center of plantar area by trained experimenters blinded to the treatment condition. For optogentic experiments, the plantar surface of the hindpaws was exposed to 5 heat trials along with green light, blue light, or no light stimulation (4 mW, 20 Hz) (*34*). For chemogenetic experiments, Lang^CreERT2^; Gq-coupled DREADD or Lang^CreERT2^; Gi-coupled DREADD mice were administered i.d. injections of 5 μL of JHU37160 dihydrochloride (25 μM, Hello Bio Inc., Princeton, NJ) 15 min before testing (*16*). The beam active and idle intensities were 38 and 5, respectively. A cutoff of time of 10 s was used to prevent excessive tissue damage.

### *In vivo* Ca^2+^ imaging from DRGs

Mice were anesthetized with isoflurane (1.5% in oxygen) and maintained at ∼37 °C using a heating pad. A midline incision was made over the lower thoracic-lumbar region, and paraspinal muscles were retracted to expose the intervertebral space between T13 and L1 vertebrae, providing access to the lumbar enlargement. A small incision was made in the intervertebral membrane and underlying dura, and a polyethylene catheter (0.2 mm diameter) was inserted caudally into the intrathecal space. AAV9.CAG.GCaMP6s.WPRE.SV40 (Addgene; a gift from Douglas Kim & GENIE Project; Addgene viral prep #100844-AAV9; RRID:Addgene_100844; 2.6 × 10^13^ vector genomes/ml (*38*)) was delivered (5 µl total volume) at a rate of 1.2 µl/min, targeting lumbar dorsal root ganglia (DRGs; ∼L4 level). The catheter was left in place for 2 min to minimize reflux, then slowly withdrawn. The incision was sutured, and mice were allowed to recover for at least 2 weeks to allow transgene expression.

For *in vivo* calcium imaging, mice were anesthetized with urethane (1.5 g/kg, intraperitoneal). The L5 DRG was surgically exposed, and the overlying dura was carefully removed. Animals were maintained on a heating pad at ∼37 °C, and the vertebral column was stabilized using a custom clamp to minimize respiratory motion artifacts. Time-series images were acquired using single-photon confocal microscopy (Leica SP5, upright configuration; Leica Microsystems) with a 10× objective (NA 0.3). Z-stack imaging (10 optical sections; 512 × 512 pixels per frame) was performed with an acquisition time of ∼7.5-8.5 s per stack. GCaMP6s fluorescence was excited with a 488 nm diode laser, and emission was collected at 500-550 nm. Sensory stimuli were applied to the ipsilateral hindpaw during imaging. Mechanical stimulation was delivered using a 1.4 g von Frey filament. Thermal stimulation was applied using heated water containing 0.1% Tween-20 (55 °C). For optogenetic activation, blue light (7 mW, 20 Hz) was delivered to the glabrous hindpaw during mechanical and thermal stimulation. For pruriceptive assays, chloroquine (5 µl; 200 µg/10 µl) was intradermally injected into the hairy skin of the hindpaw using a 30-gauge needle, and imaging was performed with or without concurrent blue light stimulation. At the end of each experiment, a high-K^+^ solution (100 mM) was directly applied to the exposed DRG to confirm neuronal viability.

Regions of interest corresponding to individual neuronal somata were manually defined. Fluorescence signals were quantified as ΔF/F₀ following background subtraction. Neurons were classified as responsive if stimulus-evoked fluorescence exceeded baseline activity by >3 standard deviations. The proportion of responsive neurons and response amplitudes were quantified and compared across experimental conditions.

### *In vivo* single-unit recording from DRGs

*In vivo* single-unit recording from DRG followed established protocol (*34*). Briefly, mice were anesthetized with urethane (1.5 g/kg, i.p.). A 3-cm-long skin incision was made aseptically at the midline of the lower back. The overlying muscles were dissected and removed from the L4-L5 spinous processes. To expose the L4 or L5 DRGs, a small dorsolateral laminectomy was performed by removing the processus accessorius and part of the processus transversus. To loosen the neuronal somata from adjacent cells, collagenase P (3 mg/ml; Sigma Millipore) was applied to the DRG, incubated for 5 min, and washed 3 times with ACSF. Collagenase application was repeated 4 times. The DRG was continually superfused with ACSF. A tungsten microelectrode was driven into the DRG by hydraulic microdrive to record extracellular single-unit activity, which was amplified and digitized using Powerlab and Spike2 software.

Units were categorized as LTMRs if they responded maximally to light touch. To estimate conduction velocity, transcutaneous electrical stimuli (0-8 mA, 2 ms) were delivered from a pulse stimulator (model 2100; A-M Systems) to receptive fields. Units were further classified by conduction velocity as follows: Aβ- (>9.0 m/s), Aδ- (1.0-9.0 m/s), and C-fibers (<1.0 m/s) (*39–41*). Each unit was retested with 1.4 g von Frey filament stimuli with and without concomitant blue light exposure (7 mW, 20 Hz).

### Tissue preparation

Skin was dissected, fixed in Zamboni fixative solution (Newcomer Supply), soaked in 30% sucrose, frozen in optimal cutting temperature compound (OCT; Tissue-Tek, Sakura Finetek), and sectioned at 40 μm using a cryostat. Free-floating sections were stored in antifreeze solution at -20°C until staining. DRG were dissected, fixed in 4% paraformaldehyde (PFA; Newcomer Supply), soaked in 30% sucrose, frozen in OCT, and sectioned at 16 μm using a cryostat. DRG sections were mounted on slides and stored at -80°C until staining.

### Immunohistochemistry

Sections were washed with PBS and then blocked for 2 h at room temperature in PBS containing 5% normal donkey or goat serum and 0.2% Triton X-100. Sections were then incubated overnight at 4°C with primary antibodies diluted in blocking buffer: NEFH (1:200; AB1989; Millipore Sigma, Burlington, MA; RRID: AB_11212727), CGRP (1:400; Bio-Rad Laboratories, Inc., Hercules, CA; RRID: AB_2290729), GFRα2 (1:800; AF429; R&D systems, Pittsburgh, PA; RRID: AB_2294621), Langerin (1:500; E-17; Santa Cruz, Dallas, TX; RRID: AB_2074213), GFP (1:500; A-11122; Invitrogen/Thermofisher, Waltham, MA; RRID: AB_221569), HA (1:500; C29F4; Cell Signal technology, Danvers, MA; RRID:AB_1549585), Semaphorin7a (1:300; bs2702R; Bioss Inc, Woburn, MA; RRID: AB_10856815), Plexin C1 (1:200; AF5375; R&D Systems; RRID:AB_2284038), biotin-XX conjugated-IB4 (1:1000; I21414; Invitrogen/Thermofisher). After washing, sections were incubated for 30 min at 37°C with corresponding secondary antibodies conjugated with Alexa Fluor 488, 555, or 647 (1:300, Jackson ImmunoResearch Inc, West Grove, PA) or with Alexa Fluor 488-conjugated Streptavidin (1:300; S11223; Invitrogen/Thermofisher) for biotinylated IB4.

For β-tubulin staining, sections were blocked with 5% goat serum and 0.2% Triton X-100 in PBS for 2 h at room temperature and then incubated with goat anti-mouse IgG (Fab fragment; 1:10; Jackson ImmunoResearch Laboratories Inc.; RRID: AB_2338476) in PBS for 1 h at room temperature. Sections were then incubated with mouse β-tubulin antibody (1:250; BioLegend Inc., San Diego, CA; RRID: AB_2313773) at 4°C for 48 h, followed by incubation with Alexa fluor 488-conjugated with goat anti-mouse IgG2a (Fcγ-specific; 1:300; Jackson ImmunoResearch Laboratories Inc.; RRID: AB_2632535) for 30 min at 37°C.

After secondary antibody incubation, sections were washed three times in PBS and mounted with antifade mounting medium containing DAPI (Vector Laboratories Inc, Newark, CA). For optical clearing, sections were incubated with DAPI (100 ng/ml) for 3 min at room temperature in the dark, mounted onto slides with 0.1-mm spacers (iSpacer; SunJin Lab), and immersed in clearing solution (RapiClear 1.49; SunJin Lab) prewarmed to 37°C. Slides were maintained at room temperature for 24 h and subsequently stored at 4°C. Imaging was performed within 1 week of clearing.

Fluorescence images were acquired using an epifluorescence microscope (Leica DM6000 B; Leica Microsystems) with a 20× objective or a confocal microscope (Leica SP5; Leica Microsystems) with a 63× objective. Confocal z-stacks were collected at 1 μm intervals with a 1 μm pinhole.

Quantification of Langerin⁺ cells and marker co-expression was performed manually by a trained observer. Spatial relationships between Langerhans cells and cutaneous nerve fibers were assessed from sequential 1 μm optical sections by a trained observer.

### Langerhans cell isolation

Skin was acutely dissected and incubated in dispase (2 U/ml) for 60-90 min at 37°C. The epidermis was separated from the dermis and further dissociated using the Epidermis Dissociation Kit (Miltenyi Biotec) according to the manufacturer’s instructions. LCs were enriched using the Epidermal Langerhans Cell MicroBead Kit (Miltenyi Biotec). Briefly, the resulting cell suspension was filtered through cell strainers. LCs were isolated by magnetic-activated cell sorting (Miltenyi Biotec).

### Immunocytochemistry of isolated Langerhans cells

Isolated LCs were fixed in 4% PFA for 20 min, applied to gelatin-coated slides, and dried on a 37 to 40°C hot plate for 20 to 30 min. Cells were then immunostained as described above, and TrueBlack Plus (Biotium, Fremont, CA) was used to quench autofluorescence.

### *In vitro* calcium imaging

DRGs were dissected from Nav1.8^cre^;Ai14 mice and enzymatically digested at 37°C for 10 min in Hanks’s balanced salt solution (HBSS; ThermoFisher) containing 20 units/ml papain (Worthington Biochemical, Lakewood, NJ) and 6.7 mg/ml L-cysteine (Sigma), followed by 10 min at 37°C in HBSS containing 3 mg/ml collagenase (Worthington Biochemical). Ganglia were then mechanically triturated with fire-polished glass pipettes. Dissociated DRG cells were pelleted, resuspended in MEM Eagle’s with Earle’s balanced salt solution containing 50 U/ml penicillin, 50 μg/ml streptomycin, 1×vitamin and 10% horse serum (all ThermoFisher), plated on glass-bottom dishes coated with poly-D-lysine (0.1 mg/ml; ThermoFisher), and cultured at 37°C in 5% CO_2_ for 16–24 hours before imaging.

LCs were isolated as described above, pelleted, resuspended in RPMI1640 (ThermoFisher) containing 1×Antibiotic-Antimycotic, 1×Sodium Pyruvate, 1×MEM Non-Essential Amino Acids Solution, 49.5 μM 2-mercaptoethanol (all ThermoFisher), 0.58 mM CaCl_2_ and 10 ng/ml TGF-β (BioLegend), and plated on glass-bottom dishes coated with recombinant E-cadherin (10 μg/ml; R&D systems). Cells were maintained at 37°C in 5% CO_2_ for 16-24 hours before imaging.

Cells were incubated in Ringer’s solution (pH7.4; 140 mM NaCl, 4 mM KCl, 2 mM CaCl_2_, 1 mM MgCl_2_, 10 mM HEPES and 4.54 mM NaOH) with 5 μM of Fluo-4 AM and 0.05% of Pluronic F-127 (Biotium) at room temperature for 30 mins. Solutions were delivered by a solenoid-controlled perfusion system (Automate Scientific). For LC experiments, histamine (100 μM), chloroquine (100 μM) and serotonin (100 μM) were applied in randomized order, followed by calcium ionophore A23187 (30 μM; Sigma). For DRG experiments, recombinant mouse semaphorin7A (0.01, 0.1, 1, 10 ng/ml; R&D Systems) was applied, followed by 144 mM potassium. Each stimulus was applied for 30 sec. In a separate set of experiments, histamine (5 μM) or chloroquine (100 μM) was applied twice, 5 min apart, with or without semaphorin7A (0.1 ng/ml), followed by 144 mM potassium. For neutralizing antibody experiments, anti-Plexin C1 antibody (10 μg/ml; MAB3887; R&D Systems; RRID:AB_10890747) or control mouse IgG (10 μg/ml; MAB0042; R&D Systems; RRID:AB_471245) were applied 30 min before imaging.

Images were obtained with an inverted microscope (Eclipse TE2000, Nikon) at excitation/emission = 480/510 nm every 3 seconds. Fluorescence intensities were normalized to baseline (F/F_0_). Cells were scored as responsive if fluorescence increased by 5 SD above the resting baseline following chemical application. Only cells responsive to high-potassium or ionophore solution were included in analysis

### Single-cell RNA sequencing

Single-cell RNA sequencing (scRNA-seq) was performed at the Hussman Institute for Human Genomics Center for Genome Technology. Enriched LCs were resuspended at ∼600 cells/μl, as determined by an automated cell counter, and loaded onto the Chromium platform (10x Genomics) using the Chromium Next GEM Single Cell 3′ Library & Gel Bead Kit v3.1. 3,386 cells were captured for library preparation, followed by sequencing on an Illumina NovaSeq X Plus platform.

### Single-cell RNA-seq data processing and quality control

scRNAseq data generated from LCs were analyzed together with six replicates from a publicly available dorsal root ganglia dataset (GSE254789). All datasets were processed using Seurat (v5). For the LC dataset, an initial filtering step was applied to retain Cd207+ cells, ensuring enrichment for bona fide LC populations prior to downstream integration. For all samples, Seurat objects were constructed without initial filtering, and standard quality control metrics were computed, including number of detected genes (nFeature_RNA), total counts (nCount_RNA), and mitochondrial read fraction (percent.mt). Cells were subsequently filtered using consistent thresholds across datasets (≥1,000 genes, ≥1,000 UMIs, and ≤10% mitochondrial reads). Quality control metrics were assessed both before and after filtering to confirm consistency across samples.

### DRG atlas and cell-type annotation

To construct a reference atlas, the six DRG replicates were integrated using canonical correlation analysis (CCA) within Seurat’s layer-based integration framework. Following normalization, identification of 2,000 variable features, scaling, and principal component analysis (PCA), datasets were integrated and embedded using UMAP. Graph-based clustering was performed across multiple resolutions, and a representative resolution (0.1) was selected for downstream analyses. Clusters corresponding to non-neuronal populations were identified based on canonical marker expression (Sox2, Ednrb, Pou4f1, Avil) and removed prior to re-clustering. Remaining cells were re-clustered and annotated using a combination of marker gene expression and prior biological knowledge. Cluster identities were consolidated into major cell-type categories, with additional subtype annotations defined based on marker co-expression.

### Mapping and label transfer of LC cells

To map LC data onto the reference atlas, we employed Seurat’s anchor-based label transfer framework. LC cells were normalized, scaled, and projected into the reference PCA space. Transfer anchors were identified between the LC query dataset and the integrated DRG reference, and cell-type labels including both cluster-level and consolidated annotations were transferred using the MapQuery function. Mapped LC cells were embedded into the reference UMAP space using the learned UMAP model, enabling visualization and comparison with reference cell populations. A combined object containing both reference and LC cells was generated for downstream analyses, with a shared low-dimensional embedding used to preserve the reference structure.

### Cell-cell communication analysis

Cell-cell communication networks were inferred using CellChat on the combined data object. Raw gene expression counts were used as input, and ligand-receptor interactions were inferred using the Mus musculus CellChat database. The standard CellChat workflow was applied, including identification of overexpressed signaling genes and interactions, computation of communication probabilities, pathway-level aggregation, and network centrality analysis. Communication networks were visualized using circular network representations, and analyses were stratified to evaluate both outgoing (LC-to-DRG) and incoming (DRG-to-LC) signaling. This approach enabled characterization of the directionality and strength of intercellular communication between LC populations and DRG-derived cell types.

## Statistical analysis

Results are presented as mean ± SEM. For comparison between two groups, a two-tailed Student’s t test was used. For comparison among more than two groups, a one-way RM ANOVA followed by Tukey’s multiple comparisons test, a one-way ANOVA followed by Dunnett’s multiple comparisons test, a two-way RM ANOVA followed by Bonferroni’s multiple comparisons test, or a two-way ANOVA followed by Tukey’s multiple comparisons test was used. Statistical significance was set at *p* < 0.05 for all tests. All statistical analyses and graphs were made using GraphPad Prism10 (GraphPad Software, San Diego, CA).

## Acknowledgments

We thank Dr. Sean Michael Boyles and Dr. William F. Hulme (Center for Genome Technology) for assistance with scRNA-seq data generation and Dr. Xinzhong Dong, Johns Hopkins University, for breeding pairs of the MrgprA3^cre^ mice.

## Author contributions

Conceptualization: TA

Methodology: HI, DP, MB, AJG, TA

Investigation: KS, HI, DP, MS, TO, JD, KS, AL, CS, ST, AJG

Visualization: AJG, TA

Funding acquisition: TA

Writing – original draft: TA

Writing – review & editing: KS, HI, DP, JD, MS, TO, KS, AL, CS, ST, MB, AJG, TA

## Figures

**Fig. S1.**
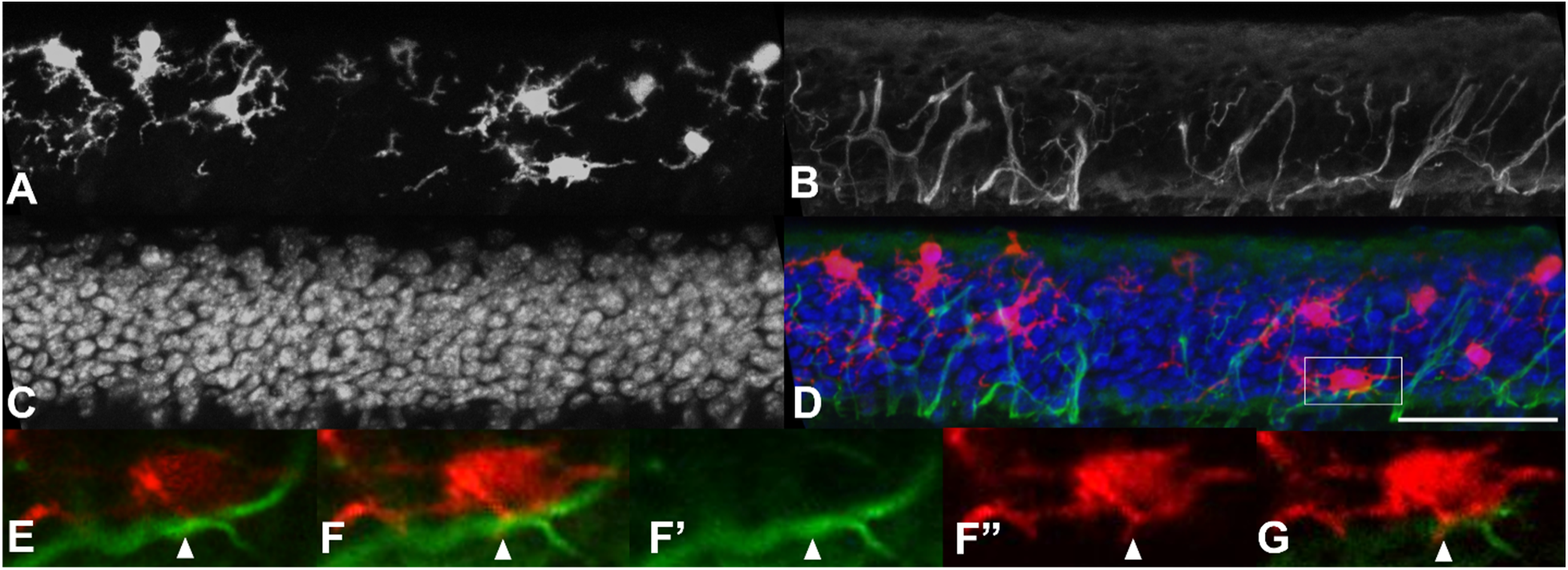
Consecutive 1-µm optical sections used to assess LC-nerve fiber proximity. **(A-D)** Example of immunohistochemical staining for LCs (tdTomato, **A**, red in **D**), peripheral nerve fibers (β-tubulin, **B**, green in **D**), and DAPI (**C**, blue in **D**) in the epidermis of the optically cleared skin. (D) Merged composite of all three channels; boxed region is magnified in (E) to (G). Scale bar = 50 µm. (**E-G)** Sequential 1-µm optical sections through the boxed region in (D), highlighting a LC colocalized with a nerve fiber (arrowhead in **F**). **F’**=green in **F**. **F”**=red in **F**.

**Figure S2.**
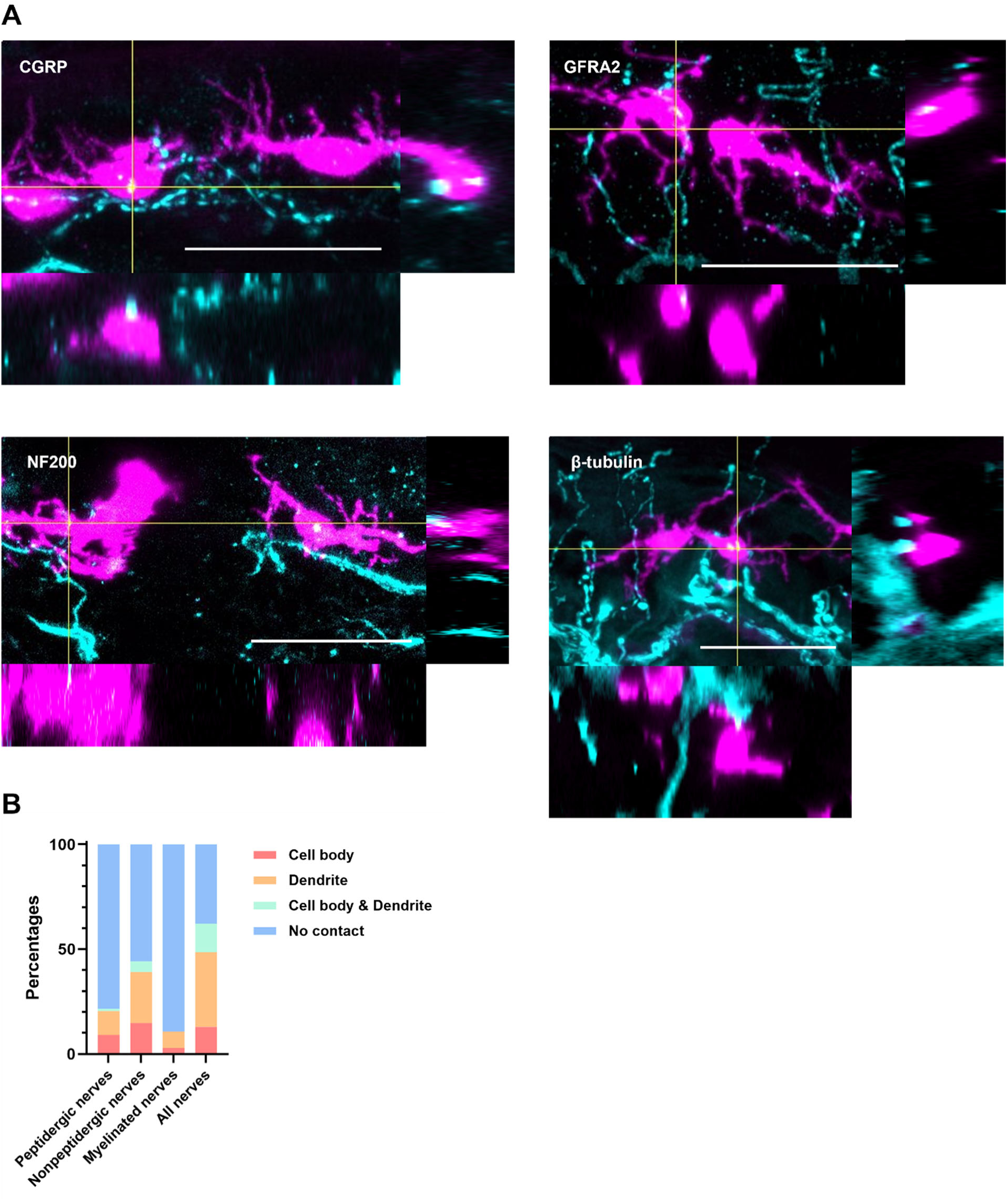
LC-nerve fiber proximity in glabrous skin. **(A-D)** Representative orthogonal projection images of optically cleared glabrous skin showing spatial proximity between LCs (magenta) and nerve fibers (cyan) labeled for peptidergic fibers (CGRP, A), non-peptidergic fibers (GFRα2, B), myelinated fibers (NF200, C), and pan-neuronal marker β-tubulin (D). Crosshairs indicate the site of LC-nerve fiber colocalization in each *xy*, *xz*, and *yz* projection. Scale bar = 30 µm. **(E)** Quantification of LC contact distribution across nerve fiber subtypes, categorized by contact site: cell body only, dendrite only, both cell body and dendrite, or no contact. *n* = 3-4 sections/mouse from 4-5 mice per marker.

**Figure S3.**
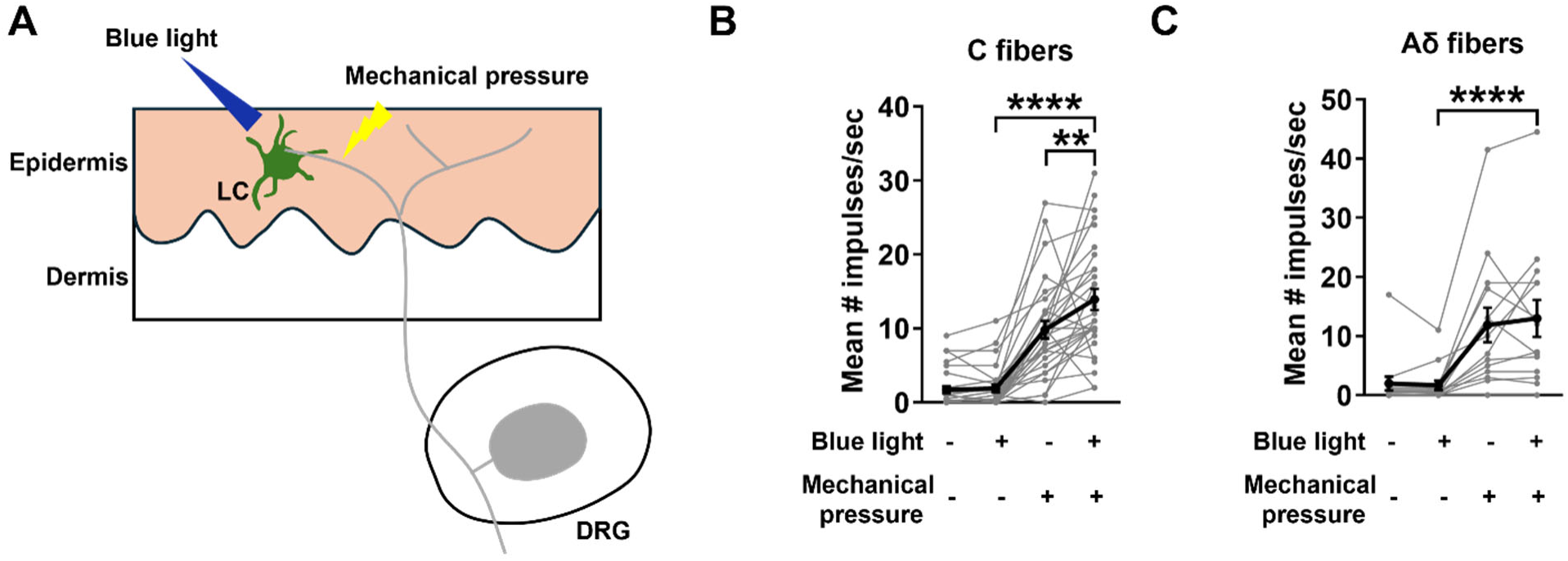
Optogenetic activation of LCs does not directly evoke neuronal firing but enhances mechanically evoked C-fiber responses. **(A)** Schematic of the *in vivo* extracellular recording preparation. A blue light stimulus (blue) activates channelrhodopsin-expressing LCs in the epidermis, while concurrent mechanical stimulation (yellow arrow) drives firing in cutaneous afferents recorded from the saphenous nerve. **(B-C)** Mean action potential firing rates in C fibers (B) and Aδ fibers (C) under four conditions: light alone, mechanical stimulation alone, combined light and mechanical stimulation, and baseline. Light stimulation alone evokes no firing in either fiber type. Combined optogenetic LC activation and mechanical stimulation significantly increases firing frequency in C fibers (B) but not Aδ fibers (C). *n* = 29 (C fibers), 14 (Aδ fibers); one-way repeated-measures ANOVA with Tukey’s post hoc test: C fibers, F(3,84) = 52.33, *p* < 0.0001; Aδ fibers, F(3,39) = 17.40, *p* < 0.0001. \*\**p* < 0.01, \*\*\*\**p* < 0.0001.

**Figure S4.**
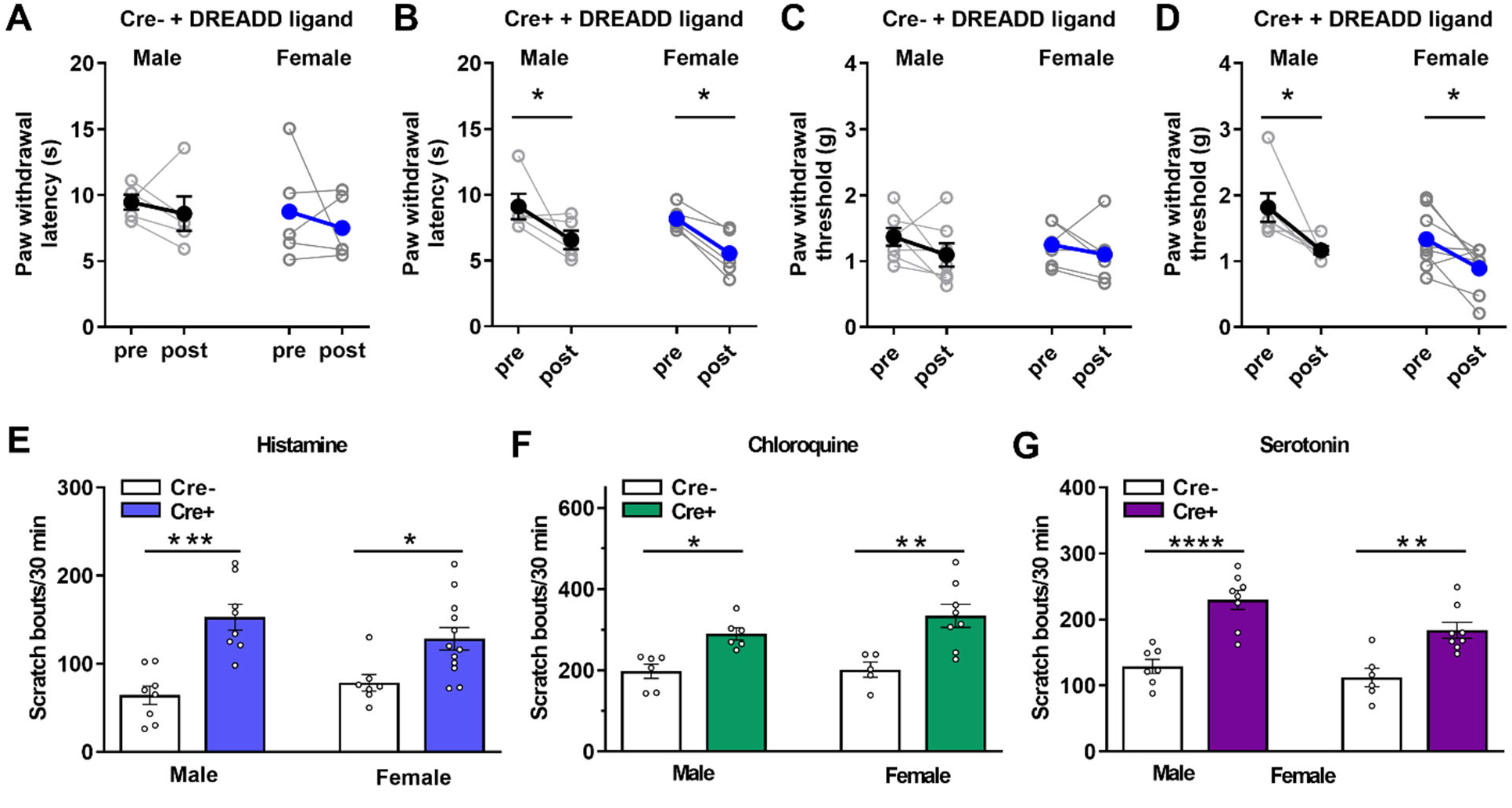
Chemogenetic LC activation enhances nociception and itch comparably in male and female mice. **(A-B)** Paw withdrawal latency to radiant heat before (pre) and after (post) injection of JHU37160 in Cre⁻ (A) and Cre⁺ (B) mice, shown separately for males and females. JHU37160 reduces thermal withdrawal latency in Cre⁺ mice of both sexes, with no effect in Cre⁻ littermates. *n* = 5 per group; two-way repeated-measures ANOVA with Fisher’s LSD test: Cre⁺, F(1,8) = 12.73, *p* = 0.0073; Cre⁻, F(1,8) = 0.778, *p* = 0.404. No significant sex differences were observed (all *p* > 0.35). **(C-D)** Mechanical paw withdrawal threshold before and after JHU37160 injection in Cre⁻ (C) and Cre⁺ (D) mice. JHU37160 reduces mechanical sensitivity in Cre⁺ mice of both sexes, with no effect in Cre⁻ controls. *n* = 6-8 per group; two-way repeated-measures ANOVA with Fisher’s LSD test: Cre⁺, F(1,12) = 14.83, *p* = 0.0023; Cre⁻, F(1,11) = 2.320, *p* = 0.156. A significant sex difference was detected only in the Cre⁺ pre-treatment baseline (*p* = 0.037); all other between-sex comparisons were not significant (*p* > 0.21). **(E-G)** Scratching bouts evoked by intradermal histamine (E), chloroquine (F), or serotonin (G) in Cre⁻ and Cre⁺ mice, shown for males and females separately. LC activation significantly increases pruritogen-evoked scratching in both sexes for all three agents. *n* = 7-12 (histamine), 5-8 (chloroquine), 6-8 (serotonin) per group; two-way ANOVA with Tukey’s multiple comparisons test: histamine, F(1,31) = 28.72, *p* < 0.0001; chloroquine, F(1,21) = 23.63, *p* < 0.0001; serotonin, F(1,25) = 43.86, *p* < 0.0001. \**p* < 0.05, \*\**p* < 0.01, \*\*\**p* < 0.001, \*\*\*\**p* < 0.0001.

**Figure S5.**
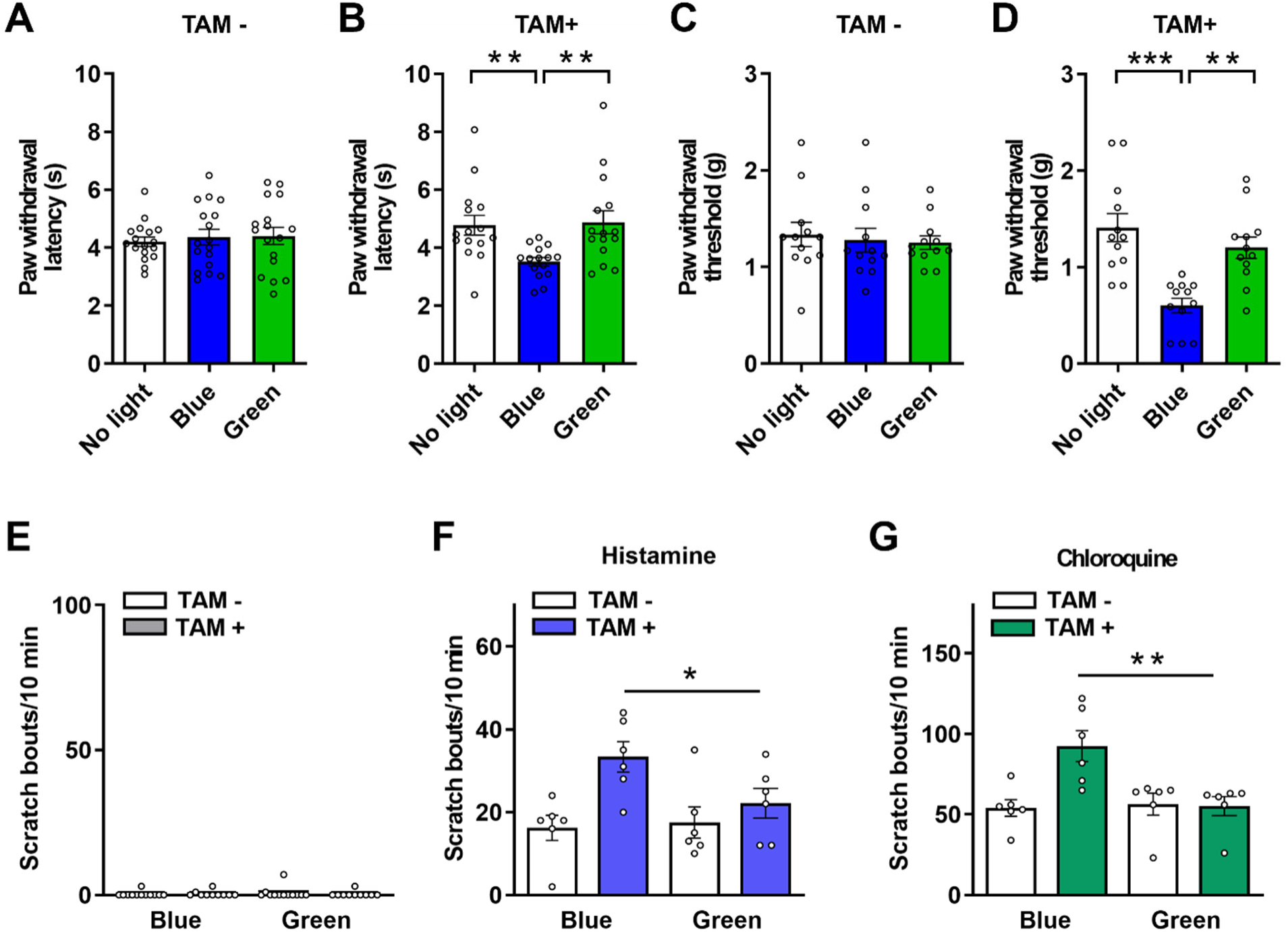
Optogenetic LC activation enhances nociceptive sensitivity and evoked scratching in tamoxifen-treated mice. **(A-B)** Paw withdrawal latency to radiant heat following blue light (channelrhodopsin activation) or green light (control) stimulation in tamoxifen-untreated (TAM⁻, A) and tamoxifen-treated (TAM⁺, B) mice. Blue light reduces thermal withdrawal latency selectively in TAM⁺ mice; green light has no effect in either group. *n* = 17 (A), 15 (B) per group; one-way repeated-measures ANOVA with Tukey’s post hoc test: TAM⁻, F(1.634, 26.14) = 0.340, *p* = 0.672; TAM⁺, F(1.856, 25.98) = 10.94, *p* = 0.0005. **(C-D)** Mechanical paw withdrawal threshold following blue or green light stimulation in TAM⁻ (C) and TAM⁺ (D) mice. Blue light reduces mechanical sensitivity only in TAM⁺ mice; green light has no effect in either group. *n* = 12 per group; one-way repeated-measures ANOVA with Tukey’s post hoc test: TAM⁻, F(1.868, 20.55) = 0.244, *p* = 0.771; TAM⁺, F(1.848, 20.33) = 16.80, *p* < 0.0001. **(E)** Optogenetic LC activation alone does not elicit spontaneous scratching in TAM⁻ or TAM⁺ mice under either light condition. *n* = 10-12 per group; two-way repeated-measures ANOVA with Bonferroni correction: F(1,20) = 0.562, *p* = 0.462. **(F-G)** Optogenetic LC activation potentiates scratching evoked by intradermal histamine (F) and chloroquine (G) in TAM⁺ but not TAM⁻ mice under blue light stimulation. *n* = 6 per group; two-way repeated-measures ANOVA with Bonferroni correction: histamine, F(1,10) = 7.733, *p* = 0.019; chloroquine, F(1,10) = 12.56, *p* = 0.005. \**p* < 0.05. \*\**p* < 0.01. \*\*\**p* < 0.001.

**Figure S6.**
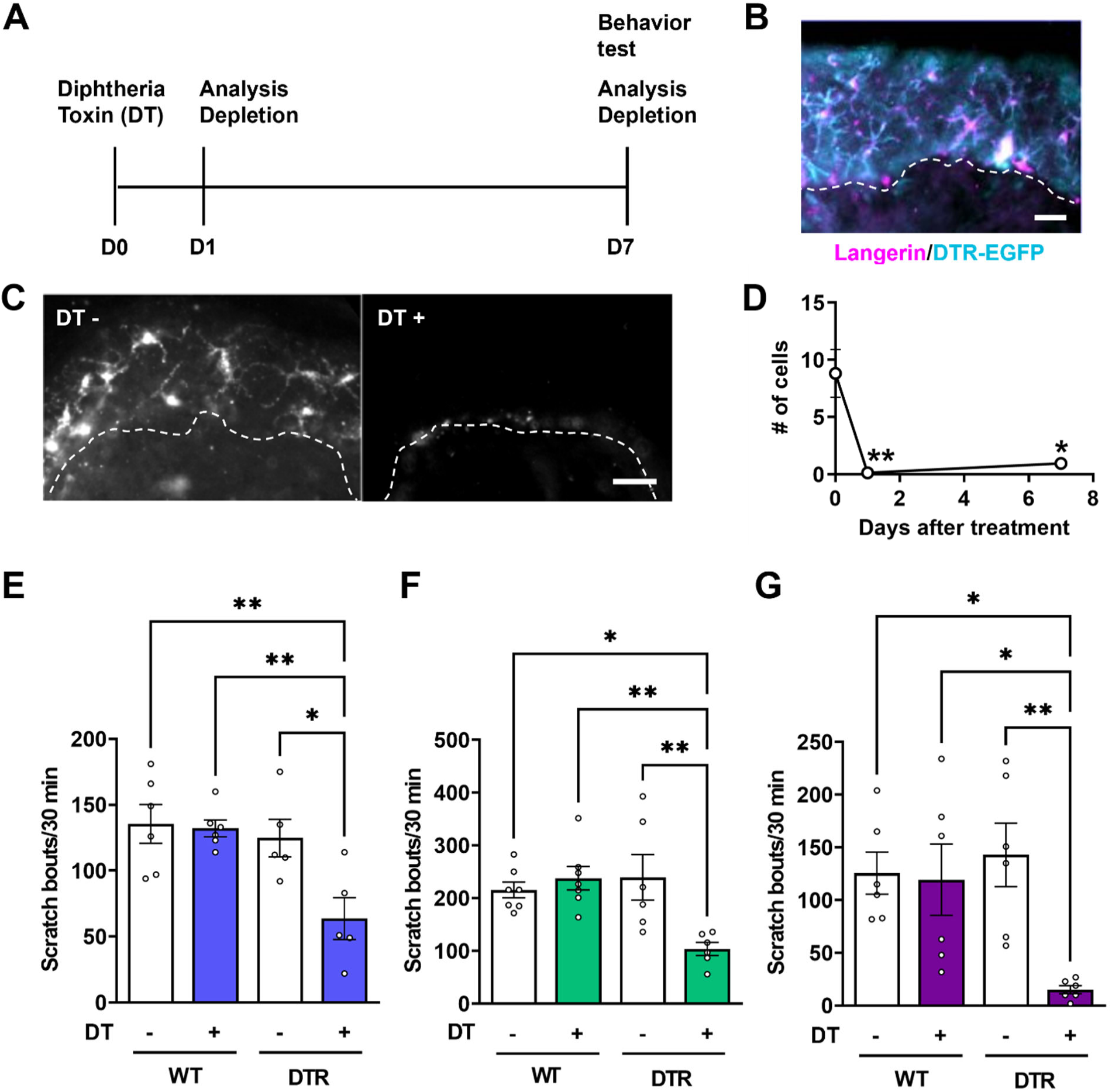
Diphtheria toxin-mediated LC depletion in Lang-DTR mice reduces pruritogen-evoked scratching. **(A)** Experimental timeline for diphtheria toxin (DT) administration and behavioral testing. **(B)** Immunostaining of Lang-DTR mouse skin showing strong co-localization of Langerin (magenta) and Langerin-DTR-EGFP (cyan), confirming selective transgene expression in LCs. *n* = 2 sections/mouse from 3 mice. Scale bar = 200 µm. **(C)** Representative immunofluorescence images of epidermal LCs (Langerin-DTR-EGFP, white) in DT-treated (DT⁺) and vehicle-treated (DT⁻) Lang-DTR mice, showing near-complete LC depletion following DT administration. Dashed line marks the dermal-epidermal junction. Scale bar = 200 µm. **(D)** Quantification of epidermal LC number at 1, 2, and 7 days after DT treatment, demonstrating sustained depletion throughout the observation period. *n* = 3 sections/mosue from 2-4 mice per time point; one-way ANOVA with Dunnett’s multiple comparisons test: F(2,6)=16.35, *p*=0.0037. **(E-G)** Scratching evoked by intradermal histamine (E), chloroquine (F), or serotonin (G) in DT-treated and vehicle-treated WT and Lang-DTR mice. DT administration selectively reduces pruritogen-evoked scratching in Lang-DTR mice but not in WT controls. *n* = 6 per group; two-way ANOVA with Tukey’s post hoc test: histamine, F(1,18) = 4.856, *p* = 0.041; chloroquine, F(1,22) = 9.810, *p* = 0.005; serotonin, F(1,20) = 6.011, *p* = 0.024. \**p* < 0.05; \*\**p* < 0.01.

**Figure S7.**
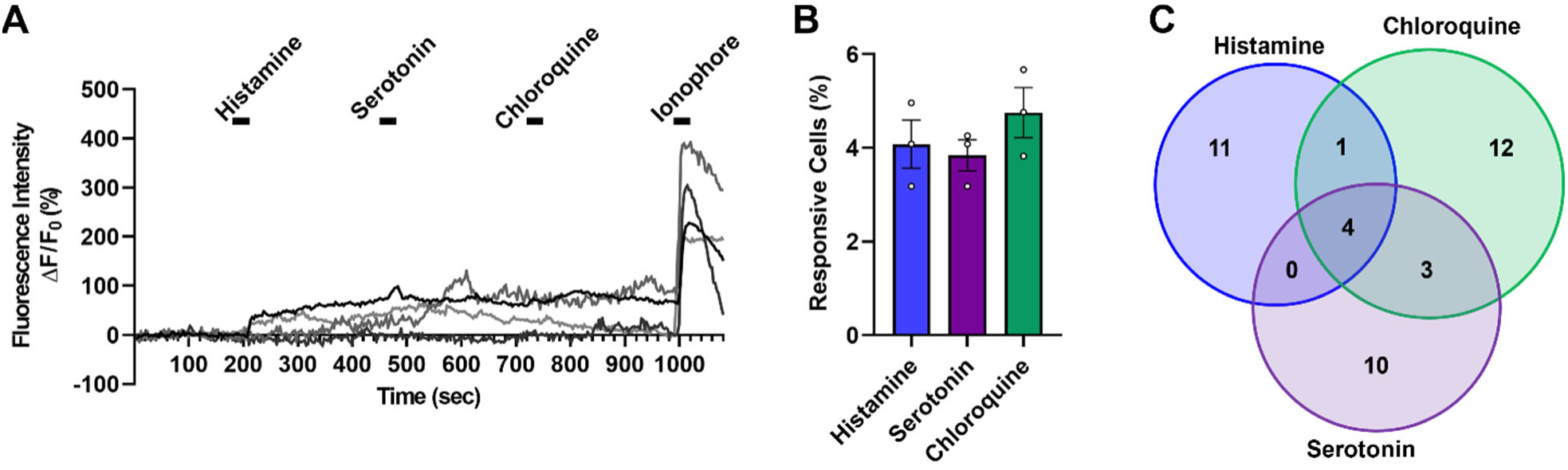
A subset of Langerhans cells directly responds to pruritogens. **(A)** Representative calcium imaging traces from cultured LCs showing fluorescence intensity (ΔF/F₀, %) over time in response to sequential application of histamine, serotonin, and chloroquine, followed by ionophore A23187 as a positive control to confirm cell viability. Individual traces (gray or black lines) illustrate heterogeneous response profiles across pruritogens. **(B)** Quantification of the percentage of LCs responding to histamine, serotonin, or chloroquine (mean ± SEM; *n* = 4 dishes/mouse from 3 mice. Open circles represent individual replicates. **(C)** Venn diagram showing the overlap among LCs responding to histamine, chloroquine, and serotonin (total responsive cells = 41/406, 10%). Response profiles were largely non-overlapping, with the majority of responsive cells reacting to only a single pruritogen (11 histamine-only, 12 chloroquine-only, 10 serotonin-only) and a minority responding to two or more agonists (1 histamine+chloroquine, 3 chloroquine+serotonin, 0 histamine+serotonin, 4 responsive to all three).

**Table S1.**
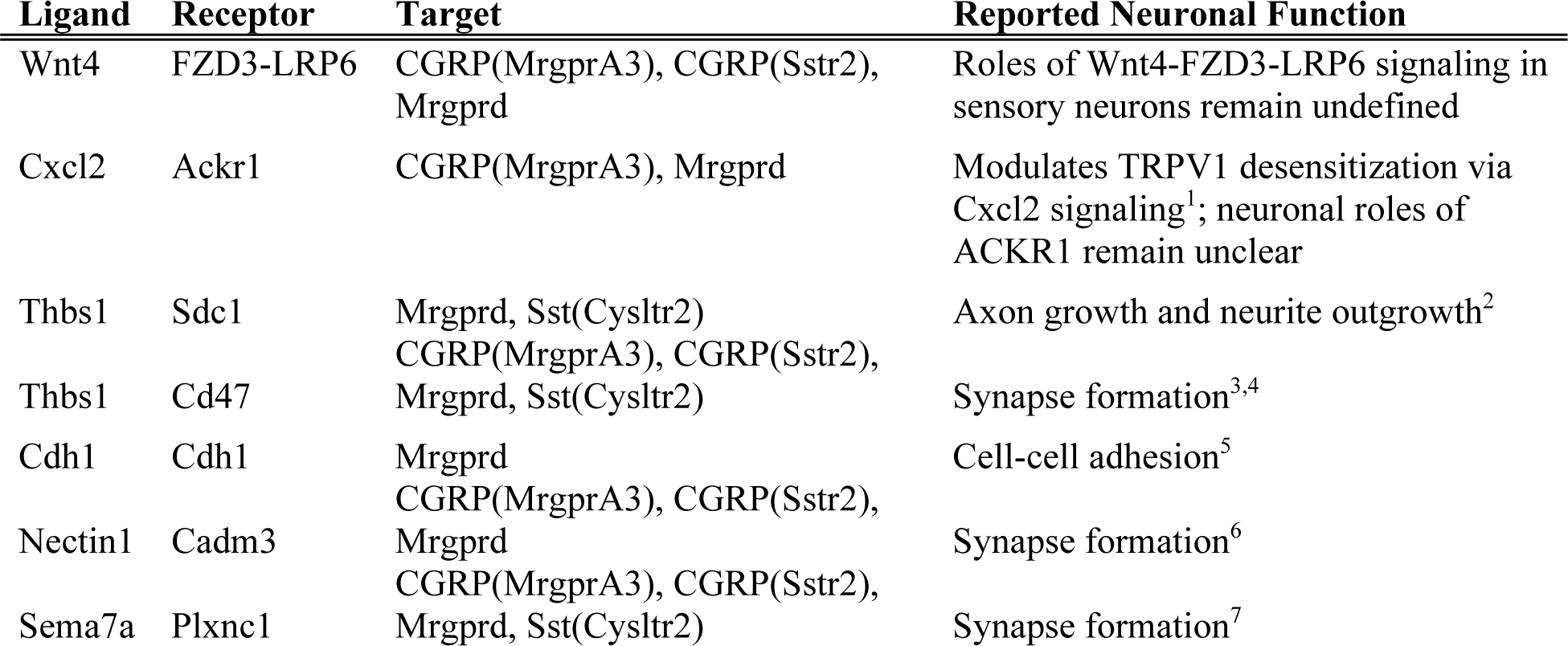
Putative ligand-receptor mediators of Langerhans cell-sensory neuron communication identified by CellChat analysis. Functions listed are based on reported roles in neuronal systems and are not restricted to sensory neurons unless indicated. **References:** ^1^Deftu, AF et al., *J Physiol Pharmacol, 2017* ^2^Bray, ER et al., *Neuron, 2019* ^3^Ding X et al., *Nat Commun, 2021* ^4^Christopherson, KS et al., *Cell, 2005* ^5^Chappuis-Flament, S et al., *JCB, 2001* ^6^Kawashima, R et al., *iScience, 2024* ^7^Inoue, N et al., *Nat Commun, 2018*

| Ligand | Receptor | Target | Reported Neuronal Function |
|---|---|---|---|
| Wnt4 | FZD3-LRP6 | CGRP(MrgprA3), CGRP(Sstr2), Mrgprd | Roles of Wnt4-FZD3-LRP6 signaling in sensory neurons remain undefined |
| Cxcl2 | Ackr1 | CGRP(MrgprA3), Mrgprd | Modulates TRPV1 desensitization via Cxcl2 signaling<sup>1</sup>; neuronal roles of ACKR1 remain unclear |
| Thbs1 | Sdc1 | Mrgprd, Sst(Cysltr2) | Axon growth and neurite outgrowth<sup>2</sup> |
| Thbs1 | Cd47 | CGRP(MrgprA3), CGRP(Sstr2), Mrgprd, Sst(Cysltr2) | Synapse formation<sup>3,4</sup> |
| Cdh1 | Cdh1 | Mrgprd | Cell-cell adhesion<sup>5</sup> |
| Nectin1 | Cadm3 | CGRP(MrgprA3), CGRP(Sstr2), Mrgprd | Synapse formation<sup>6</sup> |
| Sema7a | Plxnc1 | CGRP(MrgprA3), CGRP(Sstr2), Mrgprd, Sst(Cysltr2) | Synapse formation<sup>7</sup> |

